# A multiplexed bioluminescent reporter for sensitive and non-invasive tracking of DNA double strand break repair dynamics *in vitro* and *in vivo*

**DOI:** 10.1101/2020.03.30.015271

**Authors:** Jasper Che-Yung Chien, Elie Tabet, Kelsey Pinkham, Cintia Carla da Hora, Jason Cheng-Yu Chang, Steven Lin, Christian Elias Badr, Charles Pin-Kuang Lai

## Abstract

Tracking DNA double strand break (DSB) repair is paramount for the understanding and therapeutic development of various diseases including cancers. Herein, we describe a multiplexed bioluminescent repair reporter (BLRR) for non-invasive monitoring of DSB repair pathways in living cells and animals. The BLRR approach employs secreted *Gaussia* and *Vargula* luciferases to simultaneously detect homology-directed repair (HDR) and non-homologous end joining (NHEJ), respectively. BLRR data are consistent with next-generation sequencing results for reporting HDR (R^2^ = 0.9722) and NHEJ (R^2^ = 0.919) events. Moreover, BLRR analysis allows longitudinal tracking of HDR and NHEJ activities in cells, and enables detection of DSB repairs in xenografted tumours *in vivo.* Using the BLRR system, we observed a significant difference in the efficiency of CRISPR/Cas9-mediated editing with guide RNAs only 1-10 bp apart. Moreover, BLRR analysis detected altered dynamics for DSB repair induced by small-molecule modulators. Finally, we discovered HDR-suppressing functions of anticancer cardiac glycosides in human glioblastomas and glioma cancer stem-like cells *via* inhibition of DNA repair protein RAD51 homolog 1 (RAD51). The BLRR method provides a highly sensitive platform to simultaneously and longitudinally track HDR and NHEJ dynamics that is sufficiently versatile for elucidating the physiology and therapeutic development of DSB repair.

## INTRODUCTION

Repairing DNA damage plays a key role in maintaining genome integrity and cell viability. One DNA repair mechanism, DNA double strand break (DSB) repair, comprises two major pathways; error-prone non-homologous end joining (NHEJ) and template-dependent homology-directed repair (HDR)(1,2). The NHEJ pathway repairs DSBs by rejoining the two broken ends, which introduces random insertions or deletions at the DSB site, resulting in disruption of the gene sequence. By contrast, the HDR pathway repairs DSBs *via* homologous recombination when a donor template with a homologous sequence is available, thereby enabling insertion of desired nucleotides into the target DNA region. Importantly, the cellular preference for particular repair pathways can affect the choice of sensitizer employed in cancer treatment, as well as the efficiency of introducing therapeutic genes(3,4).

Cancer treatment often includes radiation and chemotherapy (chemoradiotherapy), which targets tumour cells by causing DNA damage, including introducing DSBs in some cases. However, this damage is recognised and often repaired by the intrinsic DNA damage response (DDR), which reduces DNA damage-induced cell death(5). Consequently, active DNA repair mechanisms can promote therapy resistance and recurrence in various tumour types. For instance, DNA repair protein RAD51 homolog 1 (RAD51) overexpression in breast and brain cancer cells can lead to increased HDR activity, resulting in resistance to chemoradiotherapy(6–8). Fortunately, small-molecule modulators of DNA repair mechanisms have since been reported to increase the efficacy of DNA-targeting therapeutics against cancers(4), and genome editing tools are being actively investigated for therapeutic and precision diagnostic applications. Meganucleases, zinc-finger nucleases (ZFNs), transcription activator-like effector nuclease (TALEN) and clustered regularly interspaced short palindromic repeat (CRISPR)-associated protein 9 (Cas9)(9) create DSBs at target DNA sites to introduce therapeutic genes by HDR, or to knockout disease-associated genes by NHEJ(10). Much effort in gene therapy development has focused on enhancing HDR over NHEJ during DSB repair to introduce functional genes, either by controlling genome editing tools, the cell cycle(11,12), optimising donor templates(13), or using small molecules to inhibit NHEJ-related proteins(14–16). However, investigating DSB repair outcomes can be time-consuming, and typically requires disruption of cells for subsequent DNA sequence analyses. This challenge has impeded high-throughput HDR optimisation for the development of cancer and gene therapies(3).

Conventional sequencing methods involve genomic DNA extraction, PCR amplification of DSB sequences, and subsequent sequence analysis methods such as Sanger sequencing and next-generation sequencing (NGS)(17). Meanwhile, mismatch cleavage nucleases such as T7 Endonuclease I (T7E1) and Surveyor nuclease have been applied to quantify insertion and deletion (indel) frequencies(18,19). However, nuclease-based methods often underestimate indel frequencies, and are unreliable when the indel frequency is over 30% or below 3%(19–22). In parallel, PCR products amplified from DSB sites can be cloned into bacterial vectors by ligation, and numerous (>48) clones must be picked for Sanger sequencing to obtain precise DSB repair results, including mutation type and indel frequency(23). In recent years, alternative strategies including tracking of indels by decomposition (TIDE) and tracking of insertions, deletions and recombination events (TIDER) have been developed(24,25). Such strategies provide a simpler analysis method for detecting indels by directly decomposing Sanger sequencing results for 500-1,500 bp PCR products of CRISPR-Cas9-edited cells. By contrast, NGS analyses of amplified PCR products provide information on the type of DSB repair, including the type and frequency of mutation sequences, as well as long mutations(9,17). NGS data are often studied using NGS analysis tools such as CRISPResso(26) to assess CRISPR-based editing results. Although NGS can detect mutation frequencies as low as 0.01%, it is costly and time-consuming, requiring days to generate results(27).

Reporter genes such as fluorescent proteins and bioluminescent luciferases are commonly used for cost-effective analysis of DSB repair results(28,29). DSB repair events can be quantified by knocking down fluorescent/bioluminescent reporter genes expressed in cells, and HDR efficiency can be measured by introducing reporter genes into target sequences. Fluorescent reporter-based methods do not require cell lysis and genomic DNA extraction, and instead use flow cytometry and/or a microplate reader for detection. However, most of these reporters are designed to reveal either HDR or NHEJ events in cells(28,30). By contrast, traffic light reporters (TLRs) developed by Certo *et al.* (2011) use an inactivated enhanced green fluorescent protein (EGFP) bearing an I-SceI site followed by a T2A peptide sequence and an out-of-frame mCherry to report HDR and NHEJ activities simultaneously(31). However, TLRs require flow cytometry analysis in order to quantitate DSB repair events, which limits their use for non-disruptive, longitudinal monitoring of DSB repair events.

Herein, we describe a non-invasive and highly sensitive bioluminescence repair reporter (BLRR) for longitudinal tracking of HDR/NHEJ both *in vitro* and *in vivo.* The BLRR method employs the naturally secreted *Gaussia* luciferase (Gluc) and *Vargula* luciferase (Vluc)(32) to enable non-disruptive observation of DSB repair activities by collecting and measuring bioluminescent data from a small amount of culture medium or blood. The BLRR assay exhibits high sensitivity and specificity for reporting HDR/NHEJ events, and results revealed a significant difference in the efficiency of CRISPR/Cas9-mediated editing with guide RNAs (gRNAs) only ~1-10 bp apart. Importantly, BLRR data are consistent with NGS results for detecting HDR events (R^2^ = 0.9722) and NHEJ events (R^2^ = 0.919). The BLRR method enables longitudinal monitoring of NHEJ/HDR activities in cultured cells and implanted tumours in mice. Using the BLRR system, we monitored altered DSB repair dynamics induced by small-molecule modulators, and subsequently revealed that anti-tumour cardiac glycosides inhibit HDR function in human glioblastomas (GBMs) and patient-derived GBM cancer stem cells (GSCs) *via* suppression of RAD51 recombinase.

## MATERIALS AND METHODS

### Molecular cloning of BLRR

To construct the BLRR, the Gluc sequence in CSCW2-Gluc-IRES-GFP was first inserted into the I-SceI cut site using 5’ and 3’ spacers at amino acid reside 104 while removing Q105 to E110, resulting in an inactive Gluc. Three silent mutations were next introduced into the nonsense Gluc at P116, S154 and G184 using a Q5 Site-Directed Mutagenesis Kit (E0554S, New England BioLabs, Ipswich, MA, USA) to remove internal stop codons. The Vluc sequence from CSCW2-Vluc-IRES-mCherry was amplified by Q5 High-Fidelity DNA Polymerase (M0491S, New England BioLabs) using primers containing a T2A peptide sequence. The PCR-amplified Vluc and nonsense Gluc sequences were cloned into *NcoI*-(R0193S, New England BioLabs) and *XbaI*-(R0145S, New England BioLabs) digested pENTR-LUC (w158-1; a kind gift from Eric Campeau & Paul Kaufman; Addgene plasmid #17473)(33) with HiFi assembly and an NEBuilder HiFi DNA Assembly Cloning Kit (E5520S, New England BioLabs) to create pENTR-BLRR. BLRR was then transferred to pLenti CMV Puro DEST (w118-1)(33) (a kind gift from Eric Campeau & Paul Kaufman; Addgene plasmid #17452) from pENTR-BLRR using Gateway LR Clonase II Enzyme mix (111791020, Invitrogen, Waltham, MA, USA), generating pDEST-BLRR.

pX330-U6-Chimeric_BB-CBh-hSpCas9 (pX330) was a gift from Feng Zhang (Addgene plasmid # 42230). To create the pX330 plasmid containing different gRNAs, 100 μM of gRNA-1, 2, 3, 4, 5 and 6-fwd and gRNA-1, 2, 3, 4, 5 and 6-rev (**Supplementary Table 1**) were mixed with 1 μl of NEB buffer2, heated to 95°C for 5 min, and cooled to 25°C (−5°C/min) to create primer dimers. These were annealed to pX330 digested with *BbsI* (R0539S, New England BioLabs). For the Gluc donor template plasmid (truncated Gluc; trGluc), trG-fwd and trG-rev (**Supplementary Table 1**) were used to amplify the Gluc sequence, which was subsequently subcloned into *NheI*- (R0131S, New England BioLabs) digested CSCW-Gluc-IRES-GFP using Gibson Assembly (E2611S, New England BioLabs). pCVL SFFV d14GFP EF1s HA.NLS.Sce(opt) was a gift from Andrew Scharenberg (Addgene plasmid # 31476).

### Cell culture

Human kidney 293T cells (293T; a gift Chien-Wen Jeff, National Tsing Hua University) were cultured in Dulbecco’s modified Eagle’s medium (DMEM; Hyclone Laboratories, SH3022.01, Logan, UT, USA) supplied with 4 mM L-glutamine, 4,500 mg/L glucose, 10% of fetal bovine serum (FBS; Hyclone, Logan, UT, USA) and 1% penicillin-streptomycin 100× solution (SV30010, Hyclone) at 37°C and 5% CO_2_ in a humidified incubator. U87-MG cells were obtained from the American Type Culture Collection (ATCC) and maintained under the same conditions. Primary GSCs used in this study were derived from a surgical specimen obtained from a GBM patient at the Massachusetts General Hospital (provided by Dr. Hiroaki Wakimoto) under appropriate Institutional Review Board approval (2005P001609). GSCs were maintained as neurospheres in DMEM/F12 medium supplemented with B27 without vitamin A (1:50; Life Technologies, Eugene, OR, USA), heparin (2 μg/mL; Sigma Aldrich, Louis, MO, USA), human recombinant EGF (20 ng/mL; ABM, Richmond, BC, Canada) and human recombinant bFGF-2 (10ng/mL; ABM). Cells were monitored for mycoplasma contamination using MycoAlert (Lonza, Basel, Switzerland). Primary cell cultures were tested monthly for mycoplasma using a PCR Mycoplasma Detection Kit (Applied Biological Materials, Richmond, BC, Canada).

### Transfection

293T or 293T cells (1×10^5^) stably expressing BLRR were seeded in 24-well plates for 24 h prior to transfection. Transfection was performed in triplicate using 0.05 mg/mL linear polyethyleneimine (PEI. molecular weight 25,000; 43896; Alfa Aesar, Heysham, Lancashire, UK) to mix 150 ng pX330-gRNA and 150 ng trGluc in 50 μl of Opti-MEM (51985091, Gibco, Waltham, MA, USA).

### Lentivirus production and generation of stable BLRR cells

For lentivirus packaging, 293T cells (1.5×10^6^) were cultured with Opti-MEM (51985091, Gibco) in 10 cm plates and co-transfected with 5 μg plasmids encoding BLRR, trGluc or SceI, 1.25 μg PMD2.G (a kind gift from Didier Trono, Addgene plasmid #12259) and 3.75 μg psPAX2 (a kind gift from Didier Trono, Addgene plasmid #12260) using PEI (43896; Alfa Aesar) in a 1:3 ratio (total DNA:PEI). At 72 h post-transfection, virus-containing medium was centrifuged at 500 × *g* for 10 min to remove cell debris, and the supernatant was filtered through a 0.45 μm pore size polyethersulfone (PES) filter (Pall, Port Washington, NY) followed by aliquotting 500 μL of filtrate per microcentrifuge tube and storage at −80°C. To generate stable BLRR cells, 293T cells (3×10^5^) were seeded in a 6-well plate overnight and cultured to 70% confluence. The medium was then replaced, supplemented with polybrene (10 μg/mL; Sigma-Aldrich), and 500 μL of lentivirus was added to the well dropwise. Cells were subsequently selected by 1 μg/mL puromycin (MDbio, Taipei, Taiwan) to generate stable BLRR cells.

### Bioluminescence BLRR assay

1 mM CTZ (Nanolight, Pinetop, AZ, USA) and 6.16 mM Vargulin (Nanolight) were diluted 1:10,000 with phosphate-buffered saline (PBS) and allowed to stabilise in the dark for 30 min at room temperature. A 200 μL volume of conditioned medium was harvested per sample and centrifuged at 500 × *g* for 3 min to collect the supernatant while removing cell debris. A 20 μL sample of supernatant was loaded per well into a 96-well white plate to measure Gluc and Vluc signals using a GloMax Discover System GM3030 (Promega, Madison, WI, USA). To measure the Gluc signal, 80 μL CTZ per well was injected using an auto-injector (GM3030, Promega) at 250 μL/s, and the signal was collected measured using a 450 nm band pass filter for 0.3 s. At 1 h after CTZ administration, the Gluc signal was remeasured to ensure that Gluc activity had diminished to background levels prior to Vluc signal detection. To measure Vluc activity, 50 μL Vargulin per well was injected at 250 μL/s, and the Vluc signal was measured with a 450 nm band pass filter for 1 s.

### Cell viability assay

Cell viability was measured after collecting conditioned medium from BLRR cells by adding 1/10 volume of alamarBlue reagent (Bio-Rad, Hercules, California, USA) to samples followed by incubation at 37°C with 5% CO_2_ for 1 h. A 100 μL volume of collected medium was used for measurement by a GloMax Discover System GM300 (Promega). Signals were collected using a 520 nm excitation filter, a 1 s integration time, and a 580-640 nm emission filter. For GBM studies, cell viability was measured using CellTiter-Glo (Promega) as recommended by the manufacturer.

### Preparation of Cas9 protein and sgRNA

Cas9 recombinant protein was expressed in *Escherichia coli* BL21 (DE3) from plasmid pMJ915 (a gift from Jennifer Doudna; Addgene # 69090) and purified as previously described (34). The purified Cas9 protein was stored at −80°C in Cas9 buffer (20 mM HEPES pH 7.5, 150 mM KCl, 10% glycerol, 1 mM β-mercaptoethanol). The sgRNAs were designed using the CRISPR design tool on the Benchling website (www.benchling.com). The sgRNAs were synthesised by *in vitro* transcription (IVT) using T7 RNA polymerase and purified by 10% denaturing urea polyacrylamide gel electrophoresis (PAGE) as described previously(12). A 1000 pmol sample of PAGE-purified sgRNA was treated with 20 U of calf intestine phosphatase (M0525L; New England BioLabs) at 37°C for 3 h to remove the 5’ phosphate group to prevent triggering innate immune responses(35). The sgRNA was then extracted with a phenol-chloroform-isoamyl alcohol mix and precipitated by isopropanol. The final sgRNA products were dissolved in sgRNA buffer (Cas9 buffer with 10 mM MgCl_2_) and stored as aliquots at −80°C. The sgRNA concentration was determined with a NanoDrop Lite instrument (Thermo Fischer Scientific, Waltham, MA, USA).

### *In vitro* cleavage assay

DNA substrates were generated using Q5 High-Fidelity DNA Polymerase (M0491S; New England BioLabs) to PCR-amplify pDEST-BLRR with TIDE-1-fwd and TIDE-1-rev (**Supplementary Table 1**) at 98°C for 30 s followed by 35 cycles at 98°C for 10 s, 64°C for 30 s, 72°C for 20 s, and a final extension at 72°C for 2 min, followed by holding at 4°C. PCR products were purified using a PCR/Gel Purification Kit (Geneaid, Taipei, Taiwan). A 0.18 μM sample of sgRNA was mixed with 0.18 μM Cas9 protein at 37°C for 5 min to form a ribonucleoprotein (RNP) mixture. 0.15 μM purified DNA products were mix with RNP mixture and incubated in 37°C for 30 min. Samples were then subjected to electrophoresis on a 1.5% Tris/Borate/EDTA (TBE) agarose gel and stained with SYBR Safe (Life Technologies) for 1 h to visualise DNA cleavage.

### TIDE and TIDER analyses

Genomic DNA was collected using a Genomic DNA Extraction Kit (Favorgen, Pingtung, Taiwan). For gRNA test samples, the BLRR sequence was amplified by Q5 Polymerase (M0491S; New England Biolabs) using primers TIDE-1-fwd and TIDE-2-rev (**Supplementary Table 1**). For small molecule test samples, the BLRR sequence was amplified with primers TIDE-2-fwd and TIDE-2-rev (**Supplementary Table 1**). In both cases, thermal cycling was performed at 98°C for 3 min followed by 30 cycles at 98°C for 10 s, 64°C for 30 s, 72°C for 20 s, and a final extension at 72°C for 2 min, followed by holding at 4°C. PCR products were separated by a 1% agarose gel, excised, and purified by a Gel Purification Kit (Geneaid). Purified samples were subsequently sequenced using either TIDE-1-fwd or TIDE-2-fwd primers, and chromatograms were analysed by TIDE (https://tide.deskgen.com/) or TIDER (https://tider.deskgen.com/).

### Animal studies and *ex vivo* blood reporter assays

Animal studies were performed in female athymic nude mice (6-8 weeks of age). These studies were conducted under the guidelines and approval of the Massachusetts General Hospital Subcommittee on Research Animal Care (MGH Animal Welfare Assurance No.: D16-00361). 293T cells were transduced with lentivirus encoding BLRR and trGluc (control) or BLRR, trGluc and I-SceI (active BLRR reporter), and implanted subcutaneously in the flanks of mice (1×10^6^ cells/mouse) separated into two groups (n = 5/group) on day 3 posttransduction with I-SceI. Tumour volume was determined by calliper measurement. Blood collection and luciferase measurement were carried out as previously described(36). Briefly, ~30 μL of blood was collected following a small incision in the tail and immediately mixed with ethylenediaminetetraacetic acid (EDTA; 10 mM) to prevent coagulation. A 5 μL sample of blood was used for Gluc and Vluc activity measurement by adding 100 μL coelenterazine (50 μg/mL; Gluc substrate) or 100 μl of vargulin (2.5 μg/mL; Vluc substrate), respectively. Photon counts were acquired for 10 s using a GloMax Discover System GM300.

### Compound treatment

A stock solution of NU7441 (Abmole, Houston, TX, USA) was made in DMSO (Sigma-Aldrich) at a final concentration of 2×10^-3^ M, and solutions of B02 (2×10^-2^ M; Abmole) and CAY10566 (CAY; 2×10^-3^ M; Cayman Chemical, Ann Arbor, Michigan, USA) were stored at −20°C. Working solutions were prepared 30 min before treating with a final DMSO concentration of 1%. BLRR cells (1×10^5^) were seeded in 24-well plates and incubated overnight for transfection with 150 ng of pX330-gRNA and 150 ng of trGluc. At 16 h post-transfection, medium was replaced with fresh medium containing either 1% DMSO (control) or the indicated concentrations of NU7441 for 1 h, then replaced with fresh medium. At 44 h post-NU7441 treatment, medium was replaced with fresh medium and incubated for 4 h prior to medium collection for BLRR assay. For B02 treatment, BLRR cells were treated with the indicated concentrations of B02 or 1% DMSO (control) for 1 h before transfection. At 44 h post-treatment, medium was replaced and cells were incubated for 4 h followed by collection of 200 μL of medium for BLRR assay. After medium collection for BLRR analysis, cells were assessed for cell viability. To test the effects of cardiac glycosides on DDR, U87-MG and GSC cells expressing BLRR/trGluc/I-SceI were treated with ouabain, digoxin or lanatoside C. U87 cells were treated at 25 and 50 nM (ouabain and digoxin) or 50 and 100 nM (lanatoside C). GSCs were treated at 12.5 and 25 nM (ouabain) or 25 and 50 nM (lanatoside C and digoxin). Gluc/Vluc activity was measured at 48 h post-treatment and expressed as fold change compared with DMSO-treated controls.

### Next-generation sequencing

Genomic DNA was extracted with a Genome Extraction Kit (Favorgen). Q5 polymerase (New England Biolabs) and primers NGS-fwd and NGS-rev (**Supplementary Table 1**) were used to amplify the gRNA target sequence at 98°C for 2 min followed by 30 cycles at 98°C for 10 s, 66°C for 30 s, 72°C for 15 s, and a final extension at 72°C for 2 min, followed by holding at 4°C. PCR products were separated on a 1% agarose gel and purified by a PCR/Gel Purification Kit (Geneaid). PCR products were analysed by Illumina Miseq 250 bp pair-end sequencing at the Genome Research Center, Academia Sinica, Taiwan. Sequencing results were analysed using the CRISPREsso web portal with average reading quality and single bp quality >30 according to the phred33 scale (26).

### Western blotting analysis

Cells were lysed in RIPA buffer (Boston Bio Products, Ashland, MA, USA) supplemented with a cocktail of protease inhibitors (5892791001, Roche, Basel, Germany) and phosphatase inhibitor (4906845001, Roche). Protein quantification was determined using the Bradford protein determination assay (Bio-Rad). A 30 μg sample of protein was loaded and resolved on a 10% NuPAGE BIS-TRIS gel (Life Technologies), then transferred to a nitrocellulose membrane (Bio-Rad) before incubation with primary antibodies. DNA-dependent protein kinase catalytic subunit (DNA-PKcs) antibody was obtained from Santa Cruz (sc-5282, Dallas, TX, USA) and anti-phosphorylated DNA-PKcs was purchased from Abcam (ab124918, Cambridge, MA, USA). Anti-RAD51 was purchased from BIOSS Antibodies (BSM-51402M, Woburn, MA, USA) and anti-β-actin was obtained from Cell Signaling Technologies (3700, Danvers, MA, USA). Gluc antibody was obtained from New England BioLabs (E8023). GAPDH antibody was obtain from Novus Biologicals (NB300-228, Centennial, Colorado, USA). Proteins were detected using SuperSignal West Pico Chemiluminescent Substrate (#34077, Thermo Fisher Scientific).

### Statistical analysis

Results are presented as mean ± standard error of the mean (SEM) unless otherwise noted. All cell culture experiments consisted of a minimum of three independent replicates which were repeated at least three times. Statistical significance was calculated using a two-tailed Student’s t-test and one-way analysis of variance (ANOVA) including comparison with the appropriate control group, followed by Tukey’s post-hoc tests. A *p*-value <0.05 was considered significant. Statistical analysis was conducted using GraphPad Prism 7 (GraphPad Software, La Jolla California USA, www.graphpad.com).

## RESULTS

### The BLRR assay non-invasively monitors NHEJ and HDR activities *in vitro*

The BLRR consists of secreted Gluc and Vluc for simultaneous monitoring of HDR and NHEJ, respectively. HDR and NHEJ activities can thus be detected by assaying each reporter activity in a small volume *(i.e.* a few μl) of conditioned medium or blood, keeping cells and animals unperturbed for subsequent molecular analyses such as sequencing and proteomics (**Figure 1A, B**). To create the BLRR system, we replaced the Q105-E110 (QGGIGE) sequence in Gluc with a 39 bp fragment containing an I-SceI endonuclease targeting site, two spacers, and a stop codon, thereby generating early translational termination and an inactive Gluc protein (**Supplementary Figure 1**). We next inserted a 2 bp frame-shifted T2A peptide sequence(37) followed by a Vluc sequence downstream of the inactive Gluc. In addition, we designed a Gluc donor template (truncated Gluc; trGluc) containing Q105-E110 but with no luciferase activity (**Supplementary Figure 2**). When DSBs occur at the I-SceI site, trGluc replaces the premature stop codon *via* HDR and triggers Gluc expression, thereby reporting HDR activity. Meanwhile, in the absence of the trGluc donor template, one of three frameshifts from NHEJ indels will correct the frameshifted T2A-Vluc sequence, causing it to become in-frame, thereby enabling subsequent Vluc expression to report NHEJ activity (**Figure 1A**). To verify BLRR function, we used two positive control constructs, BLRR-(+)NHEJ and BLRR-(+)HDR, to simulate NHEJ and HDR repair, respectively, and confirmed the specificity of BLRR signals (**Figure 1C, D**).

**Figure 1.**
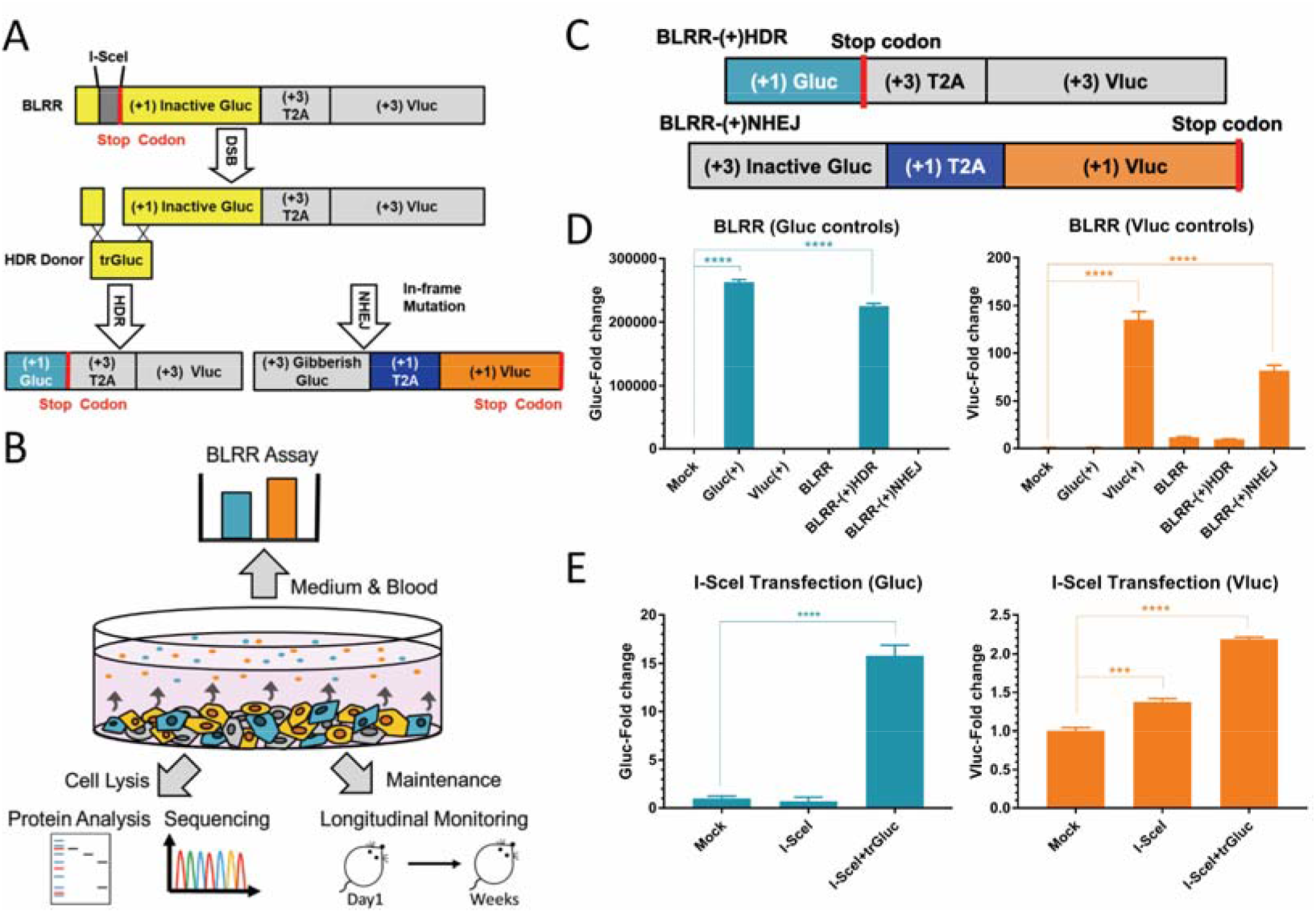
The bioluminescence DNA repair reporter (BLRR) can detect HDR and NHEJ events. (**A**) Schematic diagram and mechanism of BLRR assay detection of HDR and NHEJ repair pathways. (+) indicates the position of the reading frames with (+1) denoting the in-frame reading frame. An I-SceI meganuclease target site was inserted into the Gluc sequence followed by 2 bp frame-shifted T2A and Vluc sequences. Following DSB, NHEJ repair will generate frameshift mutations in inactive Gluc, resulting in Gibberish Gluc, and one of three frameshifts will create an in-frame T2A-Vluc sequence. When the trGluc donor template is present, HDR occurs and repairs the mutated Gluc sequence, yielding wild-type Gluc. (**B**) The BLRR system enables non-disruptive analysis of DSB repair outcomes using a small volume of medium or biofluid without disrupting cells. Cells and organisms can be further longitudinally monitored and/or collected for subsequent molecular analysis such as NGS and proteomics. (**C**) Schematic diagram of BLRR control plasmids. BLRR-(+)HDR serves as an HDR positive control by replacing the inactive Gluc sequence in BLRR with the wild-type Gluc sequence. BLRR-(+)NHEJ serves as an NHEJ positive control by replacing the I-SceI target sequence in BLRR with a +1 bp frame-shifted I-SceI target sequence to generate (+3) Gibberish Gluc, thus creating inframe T2A and Vluc. (**D**) BLRR emits Gluc and Vluc signals without signal crosstalk. 293T cells transfected with BLRR display undetectable Gluc signals and barely detectable Vluc signals. As positive controls for BLRR, 293T cells were transfected with either BLRR-(+)HDR or BLRR-(+)NHEJ, and cells exhibited robust Gluc or Vluc activity, respectively, without signal crosstalk. As positive controls for bioluminescent reporters, Gluc(+) or Vluc(+) was transfected into 293T cells to express wild-type Gluc or Vluc, respectively. A representative experiment composed of three independent experiments with three biological replicates is shown. BLRR signals were normalised against cell viability and results are shown as the fold change relative to the mock control (mean ± SEM of three biological replicates). (**E**) BLRR demonstrates I-SceI-induced DSB repair. The Vluc signal is increased in the presence of I-SceI, whereas the Gluc signal is only increased when trGluc is co-expressed. A representative experiment composed of three independent experiments with three biological replicates is shown. BLRR signals were normalised against cell viability and results are presented as the fold change relative to mock the control (mean ± SEM of three biological replicates). Significance was calculated using one-way ANOVA compared to the mock control, followed by Tukey post-hoc test (\**p* <0.05, \*\**p* <0.01, \*\*\**p* <0.001, \*\*\*\**p* <0.0001).

To examine whether the BLRR reflects endogenous DSB repair, 293T cells stably expressing BLRR (BLRR cells) were transfected with or without trGluc for 48 h to express I-SceI. Aliquots of conditioned medium were then assayed for Gluc and Vluc activities to detect HDR and NHEJ events, respectively. Importantly, the Vluc signal increased in the presence of I-SceI expression, and the Gluc signal was elevated only under co-expression of I-SceI and the trGluc donor template (**Figure 1E**).

As an alternative to I-SceI-mediated activation of BLRR, we investigated whether the BLRR can also report CRISPR/Cas9-induced DSB repair. Based on scores predicted by Benchling (http://www.benchling.com) and CHOPCHOP(38) (**Supplementary Table 2**), we selected six gRNA target sites within the I-SceI cut site to examine BLRR sensitivity for reporting gRNA editing efficiency (**Figure 2A**). We first performed *in vitro* cleavage assays with gRNAs to estimate the editing efficiency and correlate with Benchling and CHOPCHOP on-target scores, and gRNA2 yielded the lowest score, while other gRNAs exhibited a similar editing efficiency (**Figure 2B, C**). Next, BLRR cells were transfected with plasmids containing Cas9 and individual gRNAs. The BLRR assay revealed that gRNA3 exhibited the highest editing efficiency in BLRR cells, as demonstrated by elevated Vluc activity compared with the other five gRNAs, consistent with the predicted scores, except for gRNA1 (**Figure 2D** and **Supplementary Table 2**). Moreover, significant differences in Vluc activity were detected between gRNAs, suggesting that the gRNA editing efficiency varies between *in vitro* and cellular settings. No Gluc activity was observed in the absence of trGluc, indicating undetectable HDR events (**Figure 2D**). To confirm the BLRR results, we subjected the same cells to TIDE analysis(24), and demonstrated a consistent trend for indel frequency to BLRR signals in which gRNA3 yielded the highest indel frequency (**Figure 2E**). In the presence of trGluc, gRNA3 exhibited the highest Vluc and Gluc signals, demonstrating that it yielded the highest editing efficiency (**Figure 2F**). This result was further substantiated by TIDER analysis(25) on the same groups of cells, in which gRNA3 achieved the highest percentage of HDR and NHEJ events (**Figure 2G**). Interestingly, gRNA2 and gRNA4 displayed high Vluc activity but minimal Gluc activity in both BLRR and TIDER assays. Based on the these results, we selected gRNA3 to be applied with Cas9-encoding plasmids(39), hereafter referred to as px330-gRNA, for all subsequent experiments.

**Figure 2.**
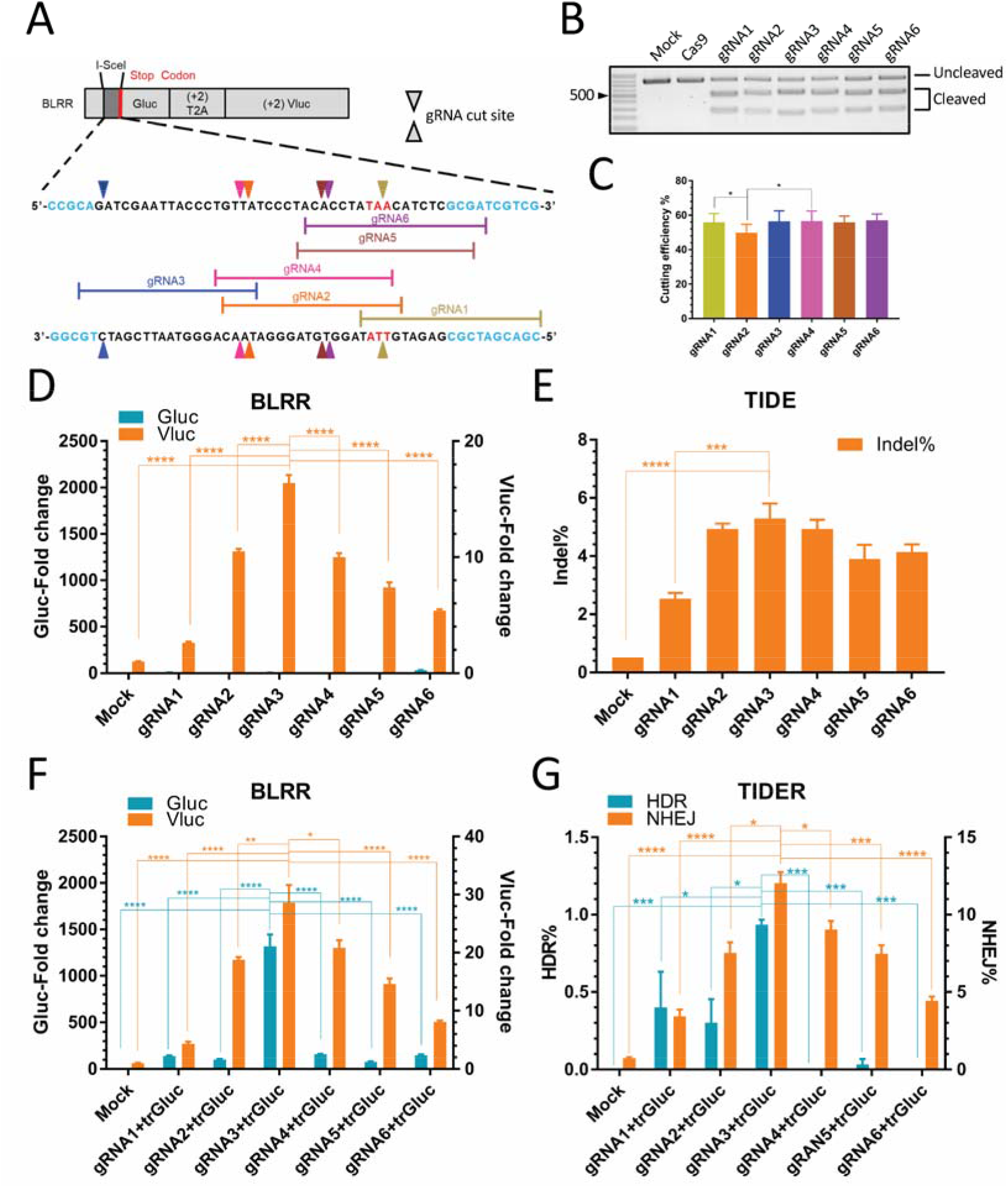
The BLRR assay can measure gRNA editing efficiency with high sensitivity. (**A**) Schematic diagram of six gRNA target sites of the BLRR. Triangles indicate gRNA cut sites; yellow, gRNA1; orange, gRNA2; dark blue, gRNA3; pink, gRNA4; brown, gRNA5; purple, gRNA6. The Gluc sequence is highlighted in light blue. A stop codon within the I-SceI insertion is highlighted in red. The *in silico* editing score of each gRNA is shown in **Supplementary Table 2**. (**B**) *In vitro* cleavage assay demonstrating different gRNA yields with varying levels of Cas9-mediated DSB. Cas9 and mock negative controls showed no detectable DSB. Representative data from three independent experiments are shown. (**C**) Statistical analysis of *in vitro* cleavage assays showing gRNA2 exhibiting the lowest editing efficiency, which corroborates the predicted scores (**Supplementary Table 2**). Data are presented as mean ± SEM of three independent experiments. (**D**) BLRR analysis showing differences in gRNA editing efficiency in cells. BLRR cells were transfected with individual Cas9-gRNA pairs, and Cas9-gRNA3 (gRNA3) and Cas9-gRNA1 (gRNA1) exhibited the highest and lowest editing efficiency, respectively. BLRR signals were normalised against cell viability and are shown as fold change relative to the mock control (mean ± SEM). A representative experiment composed of three independent experiments with three biological replicates is shown. (**E**) TIDE analysis of cells from (**D**) showing a similar trend in indel% as the BLRR assay. gRNA3 and gRNA1 exhibited the highest and lowest indel%, respectively. Data are presented as mean ± SEM of three biological replicates. (**F**) The BLRR assay can determine significant differences in HDR and NHEJ activities between different Cas9-gRNA pairs. BLRR cells were transfected with trGluc and individual Cas9-gRNA pairs at a 1:1 ratio. gRNA3 exhibited the highest Gluc and Vluc signals, whereas the other gRNAs all showed minimal Gluc activity. BLRR signals were normalised against cell viability and are presented as the fold change relative to the mock control (mean ± SEM of three biological replicates). A representative experiment composed of three independent experiments with three biological replicates is shown. (**G**) TIDER analysis of cells from (**F**) showing a similar trend for NHEJ and HDR events as reported by the BLRR assay. gRNA3 yielded the highest HDR% and NHEJ% among the gRNAs. Data are presented as mean ± SEM of three biological replicates. Significance was calculated using one-way ANOVA as indicated, followed by Tukey’s post-hoc test (\**p* <0.05, \*\**p* <0.01, \*\*\**p* <0.001, \*\*\*\**p* <0.0001).

To confirm BLRR expression under DSB repair conditions, immunoblotting analyses were performed on cell lysates of BLRR cells transfected with pX330-gRNA with or without trGluc (**Supplementary Figure 3**). Additional plasmids encoding only Cas9 (*e.g.* without gRNA; pX330), BLRR-(+)NHEJ, and BLRR-(+)HDR were used as controls. Wild-type Gluc was detected in BLRR cells transfected with pX330-gRNA+trGluc and 293T-BLRR-(+)HDR cells, confirming HDR with Gluc. Meanwhile, the end product of NHEJ, (+3) gibberish Gluc, was observed in BLRR cells transfected with pX330-gRNA and pX330-gRNA+trGluc and 293T-BLRR-(+)NHEJ. These results confirm that the BLRR system could successfully monitor HDR and NHEJ events using conditioned medium without disrupting cells.

### BLRR assay data are consistent with NGS results

To examine BLRR assay sensitivity, increasing amounts of pX330-gRNA and trGluc were introduced into BLRR cells to examine whether BLRR activity rises as DSB repair is increased. Both Gluc and Vluc signals rose when the total number of transfected plasmids increased (**Figure 3A**), demonstrating that the BLRR can quantitatively measure HDR and NHEJ. Next, we performed NGS analysis on the same cells used to generate the results shown in **Figure 3A,** and observed a similar increase in HDR and NHEJ measured by the BLRR assay (**Figure 3B**). By comparing the two assays, we verified the detection limit of Vluc to be around 14.7 ± 1.41% of NHEJ, suggesting this may be the NHEJ detection limit of BLRR (**Figure 3B,** 90+90 ng). By contrast, the Gluc signal has a detection limit of 1.23 ± 0.32% of HDR (**Figure 3B,** 60+60 ng), indicating that the BLRR system is more sensitive for detecting HDR than NHEJ. Notably, we observed a robust correlation between BLRR signals and NGS results; the coefficient of determination (R^2^) between the Gluc signal and HDR% was 0.9722 (**Figure 3C**) and the R^2^ value between the Vluc signal and NHEJ% was 0.919 (**Figure 3D**). To further validate BLRR sensitivity for reporting the type and frequency of DSB repair, an increasing amount of trGluc combined with a fixed quantity of pX330-gRNA were transfected into BLRR cells. BLRR analysis showed that the Gluc signal rose as trGluc was increased, indicating elevated HDR events (**Figure 3E**). Concurrently, NGS analysis of the same cells used to generate the results shown in **Figure 3E** demonstrated an increase in HDR events (**Figure 3F**). Meanwhile, an increase in HDR did not result in a decrease in NHEJ, as observed by both BLRR and NGS analyses. A linear relationship was observed between BLRR and NGS analyses (**Figure 3G, H**) with R^2^ = 0.9217 between HDR% and Gluc, and R^2^ = 0.7512 between NHEJ% and Vluc. Although HDR and NHEJ activities are often considered to be inversely correlated, Richardson *et al.* demonstrated an increase in error-prone repair outcomes, in addition to HDR elevation, when single- and doublestranded HDR donor DNAs were present(40,41). Our current findings concur with this observation; the introduction of trGluc donor DNA increased both HDR and NHEJ activities (**Figure 3E, F**), even though the HDR donor DNA was introduced *via* plasmids in our study. A subsequent investigation by the same group revealed that non-homologous single- and double-stranded DNA significantly stimulates Cas9- mediated gene disruption in the absence of HDR(41). Furthermore, we transfected BLRR cells with a fixed amount of trGluc and increasing quantities of pX330-gRNA, and the results demonstrated an elevation in the Vluc signal with increasing NHEJ events, while Gluc and HDR events remained relatively unchanged (**Supplementary Figure 4**). Taken together, the BLRR method reports DNA DSB repair outcomes with high specificity and sensitivity, as corroborated by concurrent NGS analysis.

**Figure 3.**
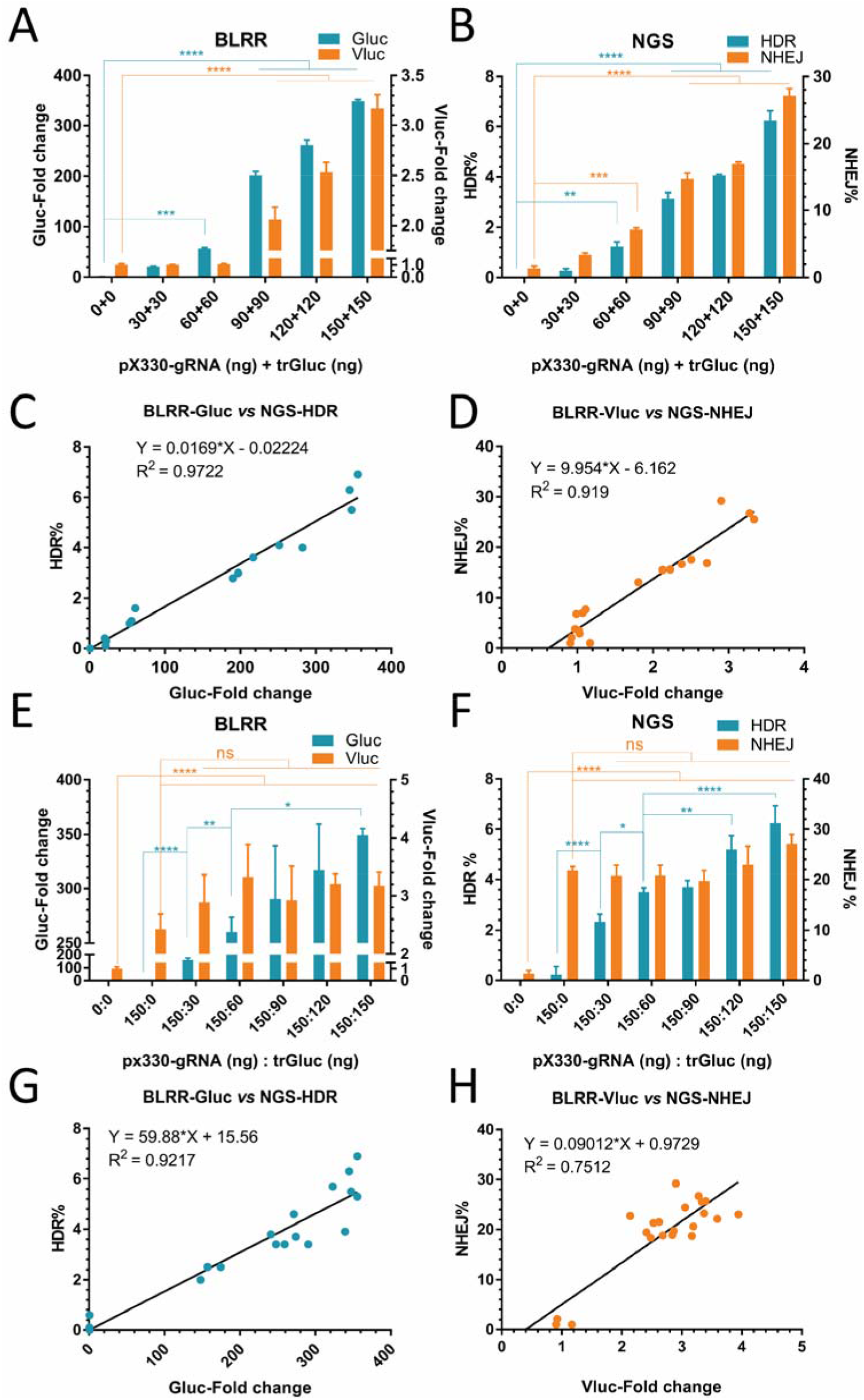
BLRR assay data are consistent with NGS results. (**A**) BLRR reporter activities increase with increasing HDR/NHEJ events. Both Gluc and Vluc signals increase as the amount of transfected pX330-gRNA and trGluc is increased. Gluc and Vluc signals exhibit significant differences at 60+60 ng and 90+90 ng compared with 0+0 ng controls. BLRR signals are normalised against cell viability and shown as the fold change relative to the 0+0 group (mean ± SEM of three biological replicates). Representative data from three independent experiments are shown. (**B**) NGS analysis of cells from (A) showing a consistent trend in HDR% and NHEJ% to those of the BLRR assay. NGS analysis of (A) showing that HDR and NHEJ events increase as the amount of transfected pX330-gRNA and trGluc is increased. Data are presented as mean ± SEM of three biological replicates. (**C**) BLRR assay Gluc values are strongly correlated (R^2^ = 0.9722) with NGS-detected HDR events. (**D**) BLRR assay Vluc values are strongly correlated (R^2^ = 0.919) with NGS-detected NHEJ events. (**E**) BLRR assay results showing that the Gluc signal is increased as the amount of transfected trGluc donor template is increased. The Vluc signal remains similar when different amounts of trGluc are applied and pX330-gRNA remains constant. BLRR signals are normalised against cell viability, and results are shown as the fold change relative to the 0+0 group (mean ± SEM of three biological replicates). Representative data for three independent experiments are shown. (**F**) NGS analysis of cells from (**E**) showing an increase in HDR events when the amount of trGluc is increased, as reported by the BLRR assay. NGS analysis of (**E**) showing an increase in HDR as the amount of trGluc is increased while NHEJ remains unaffected, corroborating the BLRR assay results. Data are presented as mean ± SEM of three biological replicates. (**G**) BLRR assay Gluc values are strongly correlated (R^2^ = 0.9217) with NGS results showing an increase in HDR events. (**H**) BLRR assay Vluc values and NGS-NHEJ (%) are strongly correlated (R^2^ = 0.7512). Significance was calculated using one-way ANOVA compared to 0+0 ng controls or as indicated followed by Tukey’s post-hoc test (\**p* <0.05, \*\**p* <0.01, \*\*\**p* <0.001, \*\*\*\**p* <0.0001).

### Longitudinal tracking of DSB repair dynamics *in vitro* and *in vivo*

Since the BLRR system employs secreted luciferases, we anticipated that it may be able to longitudinally track DSB repair events. To test this capability, we transfected BLRR cells with pX330-gRNA with or without trGluc, and measured luciferase activities using conditioned media collected at different time points (**Figure 4A**). BLRR assays showed a significant increase in Vluc and Gluc signals at 30 h post-transfection in the px330-gRNA+trGluc group (**Figure 4B, C**), and the Gluc signal reached a plateau at 48 h. Interestingly, Vluc activity displayed a slight decline at 60 h, which may be a result of cell death from the prolonged culturing time, as well as the shorter half-life of Vluc (50 h)(42) compared with that of Gluc (~6 days)(36). To validate the longitudinality of the BLRR assay, we performed NGS analysis on cells prepared in parallel with samples collected at different time points, and observed a similar increasing trend for HDR% (**Figure 4D**) and NHEJ% (**Figure 4E**). Interestingly, NGS detected increases at 24 h, 6 h earlier than the elevations observed by the BLRR assay at 30 h. The moderate difference between the two assays is likely attributed to the time required by cells to translate luciferase mRNA into enzyme following DSB repair. These results demonstrate that the BLRR can non-invasively and longitudinally monitor genome editing events.

**Figure 4.**
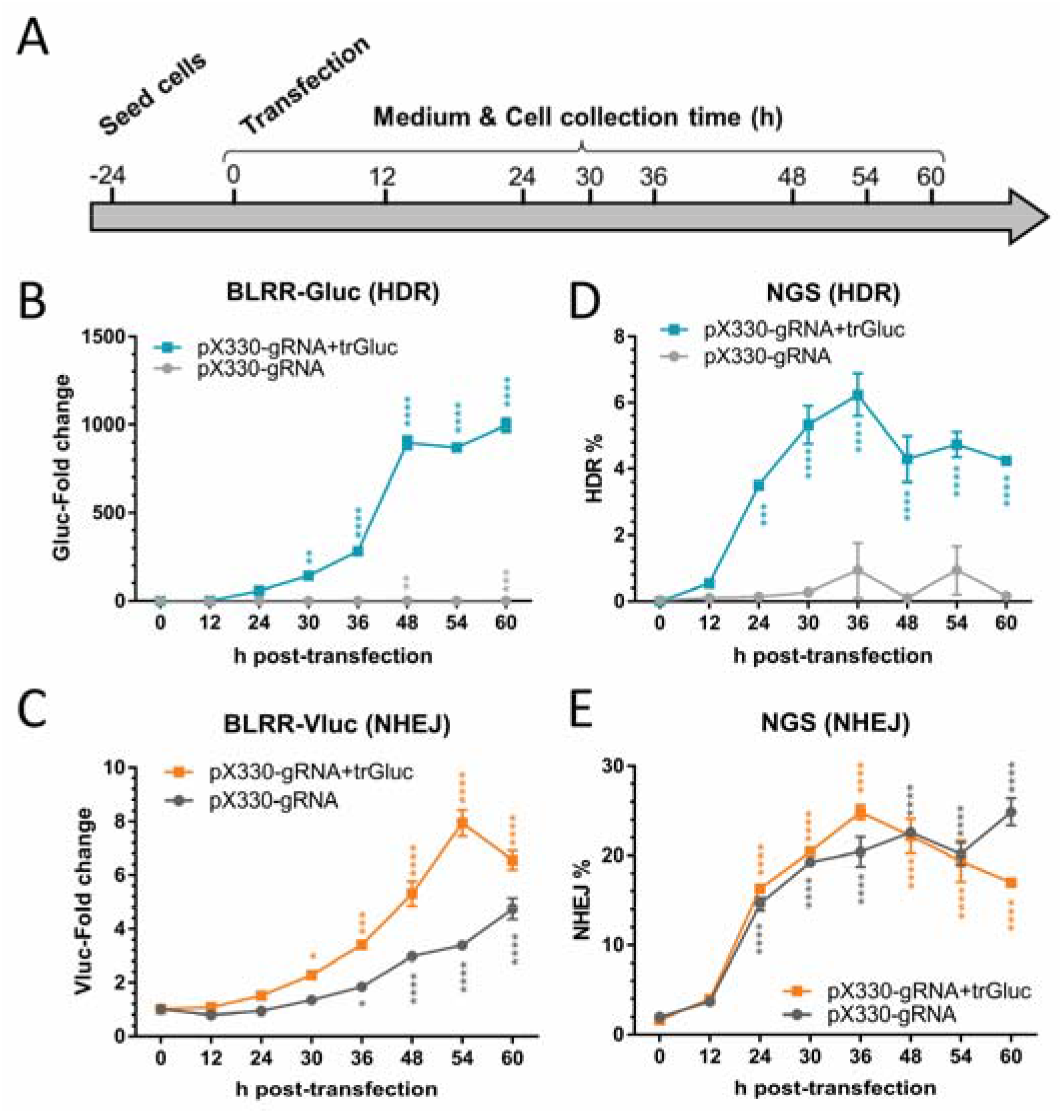
BLRR enables longitudinal tracking of HDR and NHEJ events. (**A**) Schematic diagram of longitudinal monitoring of DSB repair events. BLRR cells were transfected with or without trGluc donor template, and cells and media were collected at different time points for BLRR and NGS analyses. (**B, C**) BLRR longitudinally and simultaneously monitors HDR (**B**) and NHEJ (**C**) events. BLRR cells were transfected with either px330-gRNA+trGluc or pX330-gRNA (negative control for HDR), and both Gluc and Vluc signals showed a significant increase at 30 h compared to 0 h post-transfection. BLRR signals are shown as the fold change relative to 0 h (mean ± SEM of three biological replicates). Representative data for three independent experiments are shown. (**D, E**) NGS analysis of cells from (**B, C**) showing a similar increasing trend in HDR (**D**) and NHEJ (**E**) events, with a significant difference from 24 h post-transfection. Data are presented as mean ± SEM of three biological replicates. Significance was calculated using one-way ANOVA compared with 0 h followed by Tukey’s post-hoc test (\**p* <0.05, \*\**p* <0.01, \*\*\**p* <0.001, \*\*\*\**p* <0.0001).

Although several DDR reporters have been established, their applications have been largely restricted to cell culture models. Hence, we tested whether the BLRR could detect HDR/NHEJ in small animal models through *ex vivo* monitoring of Gluc and Vluc activities in blood samples (**Figure 5A**). We stably transfected 293T cells with BLRR+trGluc+I-SceI (active BLRR reporter) or BLRR+trGluc (negative control), and subcutaneously implanted the resulting cells in the flanks of nude mice. As the tumour size increased (**Supplementary Figure 5**), an increase in Gluc (HDR) and Vluc (NHEJ) activities was observed starting on Day 21 post-implantation in mice bearing 293T-BLRR+trGluc+I-SceI tumours, and signals increased significantly over time (**Figure 5B, C**). By contrast, low BLRR signals were detected in the 293T-BLRR+trGluc control group. The capability of the BLRR assay to longitudinally track DSB repair *in vitro* and *in vivo* will be advantageous for experiments requiring continuous monitoring of DSB repair events, as well as studies that require further molecular analysis of cells following DSB repair.

**Figure 5.**
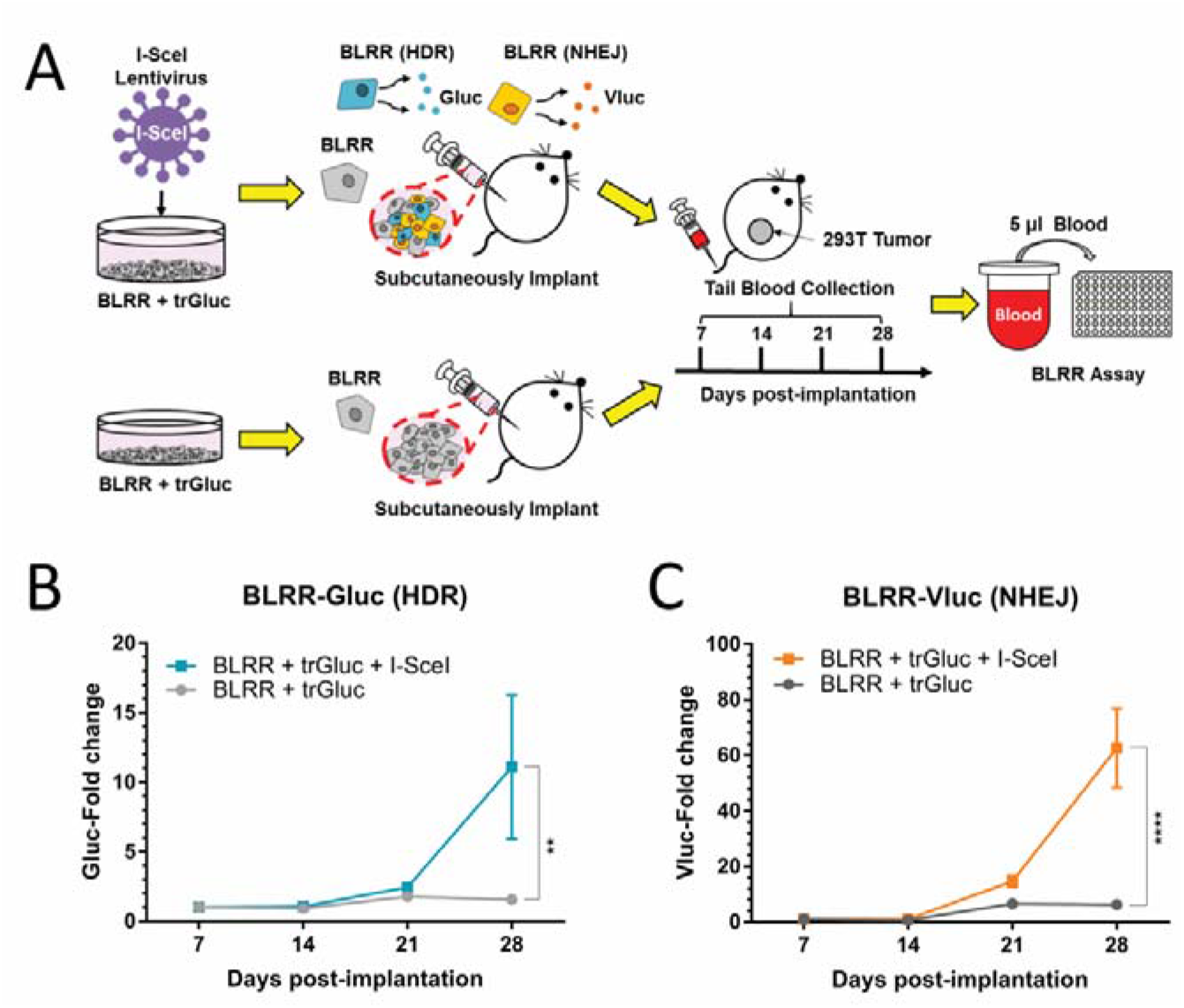
*In vivo* monitoring of HDR and NHEJ events. (**A**) Schematic diagram of longitudinal BLRR assays *in vivo.* BLRR cells were sequentially transduced to express trGluc with or without I-SceI (negative control), and subcutaneously implanted into the flanks of nude mice to facilitate tumour formation. Blood sample were collected every 7 days and Gluc and Vluc activities were measured. (**B, C**) BLRR assays of blood samples revealing an increase in Gluc (HDR) and Vluc (NHEJ) activities over time as the tumour develops (**Supplementary Figure 5**). Mice with tumours expressing BLRR+trGluc+I-SceI showed a marked increase in Gluc (**B**) and Vluc (**C**) activities compared with the BLRR+trGluc control group. BLRR signals are shown as the fold change relative to day 7 (presented as mean ± SEM of three mice). Significance was calculated by two-way ANOVA as indicated, followed by Śídák’s multiple comparisons test (\*\**p* <0.01, \*\*\*\**p* <0.0001).

### The BLRR can measure HDR and NHEJ dynamics induced by small-molecule modulators

Small-molecule compounds have been used to modulate DSB repair and enhance gene editing and therapeutic efficiencies(43). To investigate whether BLRR can effectively monitor the effects of small-molecule compounds on DSB repair, we treated BLRR cells with an HDR enhancer (NU7441) or an inhibitor (B02) and assessed HDR/NHEJ dynamics by BLRR assay. NU7441 inhibits DNA-dependent protein kinase catalytic subunits to increase HDR(44), whereas B02 inhibits RAD51 recombinase to impede HDR(45). Following NU7441 treatment, the Gluc signal increased as the Vluc signal decreased in a dose-dependent manner (**Figure 6A**). The BLRR ratio (Gluc activity divided by Vluc activity) exhibited a dose-dependent increase, suggesting that it can be applied to assess the dynamics between HDR and NHEJ events (**Figure 6B**). The same cells were further analysed by TIDER assay (**Supplementary Figure 6A**), and the value of HDR%/NHEJ% was strongly correlated with the BLRR ratio (R^2^ = 0.9594; **Figure 6C** and **Supplementary Figure 6B**). To support the BLRR results, we also examined the expression levels of key components in HDR and NHEJ pathways, namely RAD51 and phosphorylated DNA-Pkcs, and observed a dose-dependent decrease in the percentage of phosphorylated DNA-PKcs (**Supplementary Figure 6C**). By contrast, treatment with B02 resulted in a dose-dependent decline in Gluc activity in BLRR cells (**Figure 6D**). Although Vluc activity also decreased with an increasing dose of B02, the BLRR ratio showed a dose-dependent decrease, suggesting that HDR was suppressed by B02 (**Figure 6E**). TIDER analysis corroborated the BLRR assay findings, and revealed a correlation between the BLRR ratio and HDR%/NHEJ% (R^2^ =0.7411; **Figure 6F** and **Supplementary Figure 7A, B**). In addition, we observed reduced DNA-PKcs expression following B02 treatment, which likely resulted in the decreased Vluc signals, especially at higher dosages (**Supplementary Figure 7C**). These results indicate that BLRR signals and the BLRR ratio can be applied to investigate the effect of small molecules or other modalities in modulating DSB repair, which is of relevance to high-throughput screening and preclinical studies.

**Figure 6.**
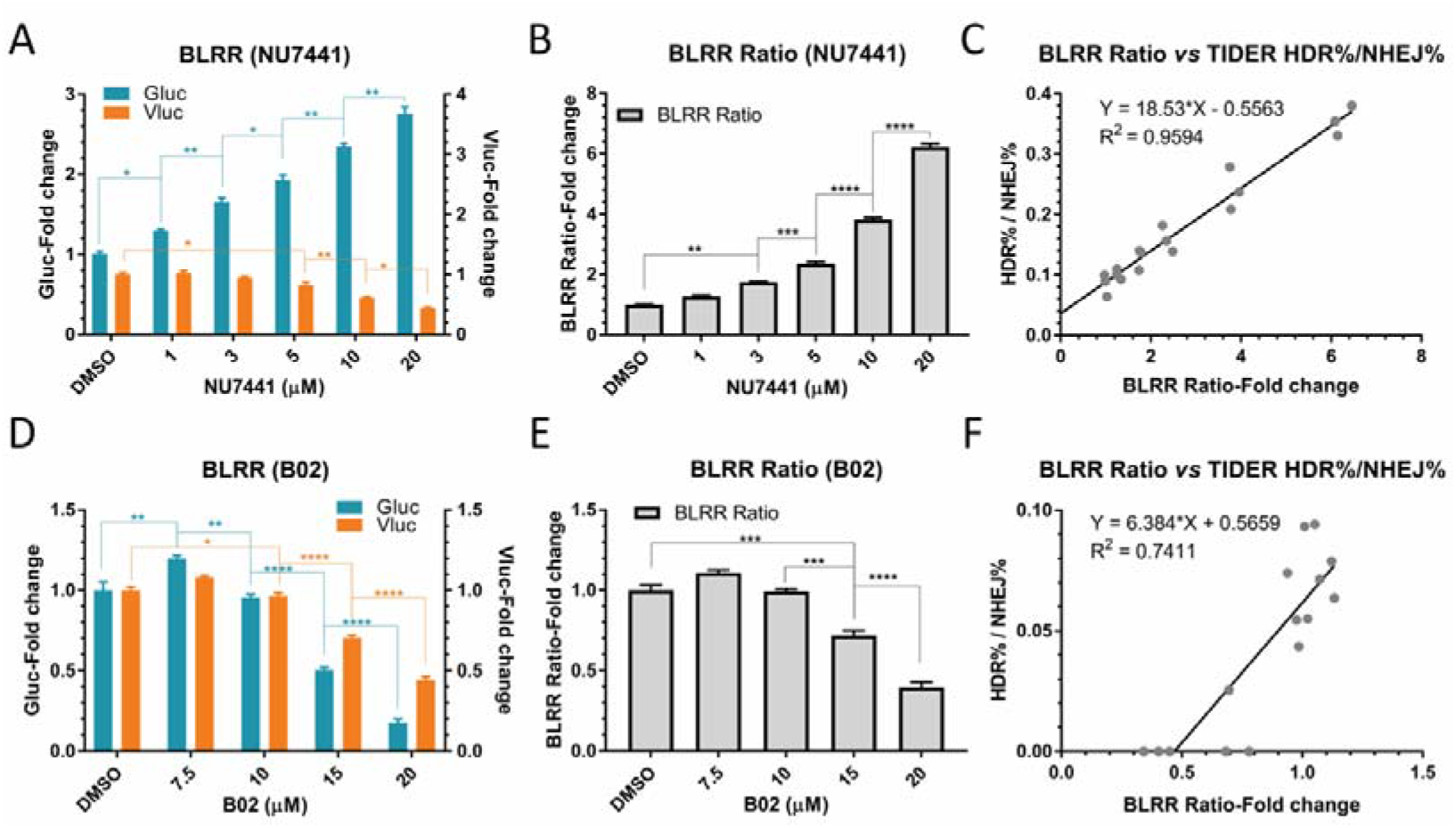
The BLRR assay can detect altered dynamics for DSB repair induced by small-molecule modulators. (**A**) BLRR activity reveals dose-dependent HDR-enhancing and NHEJ-suppressing effects of NU7441. BLRR cells were transfected with pX330-gRNA and trGluc and treated with NU7441. BLRR analysis of conditioned media revealed an increase in Gluc signal and a decrease in Vluc signal with increasing dosage of NU7441. BLRR signals were normalised against cell viability and results are shown as the fold change relative to DMSO-treated controls (mean ± SEM of three biological replicates). Representative data for three independent experiments are shown. (**B**) BLRR ratio displaying a dose-dependent increase in HDR events for NU7441. The BLRR ratio is Gluc activity divided by Vluc activity, normalised against DMSO-treated controls (mean ± SEM of three biological replicates). (**C**) TIDER analysis of **A** (**Supplementary Figure 6A, B**) showing a strong linear correlation between BLRR ratio and HDR%/NHEJ% (R^2^ = 0.9594. (**D**) The BLRR assay reveals dose-dependent HDR suppression by B02. BLRR cells were treated with B02 prior to transfection with pX330-gRNA and trGluc. BLRR analysis of conditioned media demonstrated a significant reduction in Gluc and NHEJ signals starting at 15 μM compared with the DMSO-treated control. BLRR signals were normalised against cell viability and results are shown as the fold change relative to DMSO-treated controls (mean ± SEM of three biological replicates). Representative data for three independent experiments are shown). (**E**) BLRR ratio showing a dose-dependent suppression of HDR by B02. The BLRR ratio is shown as the fold change relative to DMSO controls (mean ± SEM of three biological replicates). (**F**) TIDER analysis of **D** (**Supplementary Figure 7A, B**) reveals a linear correlation between BLRR ratio and HDR%/NHEJ% (R^2^ = 0.7411). Significance was calculated using one-way ANOVA compared with the DMSO group, followed by Tukey’s post-hoc test (\**p* <0.05, \*\**p* <0.01, \*\*\**p* <0.001, \*\*\*\**p* <0.0001).

### The BLRR assay reveals HDR-suppressing effects of cardiac glycosides in GSCs and GBM cells

Genomic instability and enhanced DNA repair are defining features of tumour cells(46). In fact, upregulation of DDR contributes to increased therapeutic resistance in stem-like tumour populations(7,47,48). Therefore, we tested whether BLRR can detect modulated DSB repair events in patient-derived GBM cancer stem cells (GSCs) (**Figure 7A**). As a positive control for BLRR detection of HDR and NHEJ activities, GSCs were transfected to co-express BLRR, trGluc and I-SceI, and a marked increase in Gluc activity (400-fold) was observed (**Supplementary Figure 8**). By contrast, only Vluc activity could be readily detected following co-expression of BLRR and I-SceI. Background Gluc and Vluc signals were detected in BLRR+trGluc and mock controls. These results indicate that the BLRR reports NHEJ and HDR events in GSCs with high specificity.

**Figure 7.**
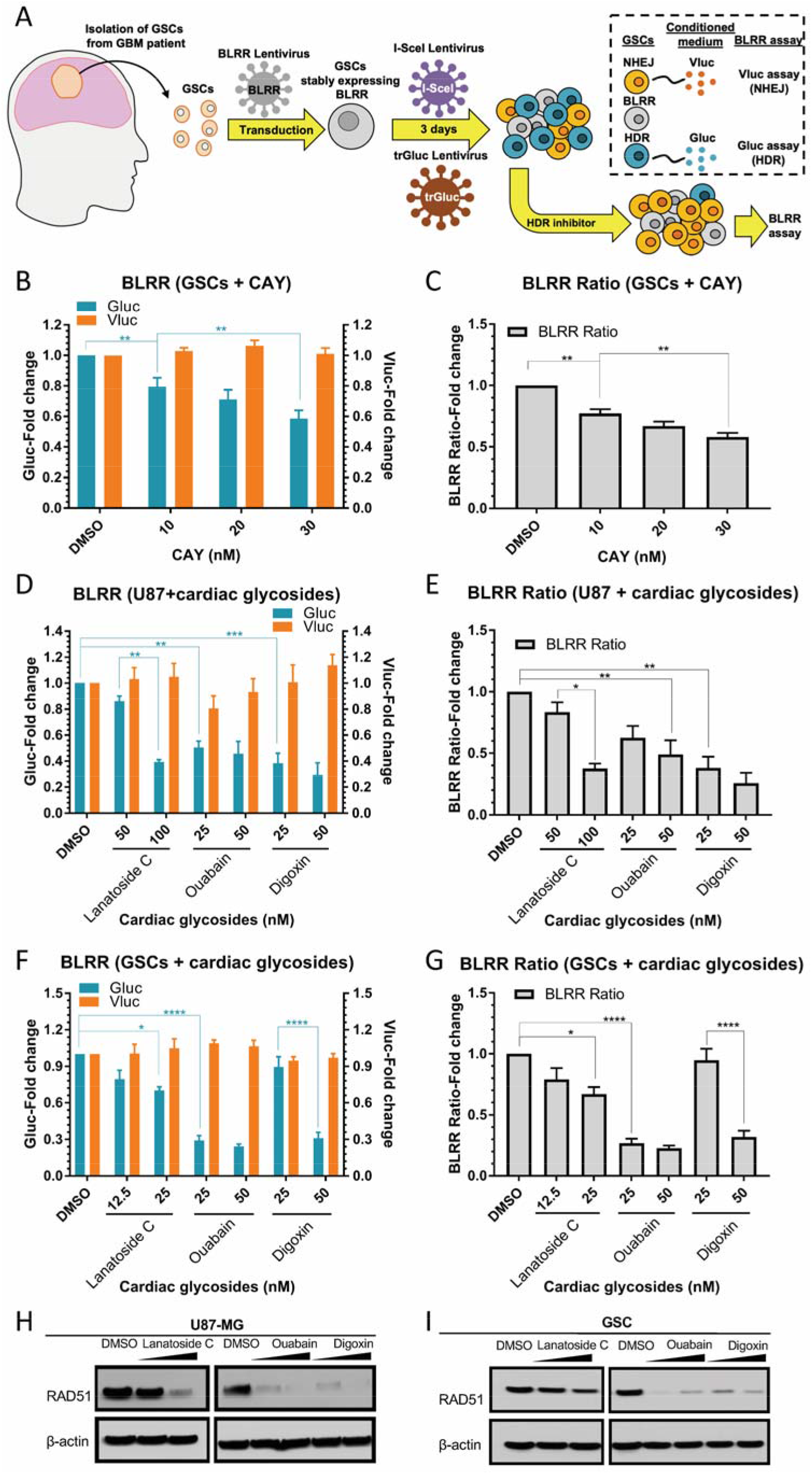
The BLRR assay reveals HDR-suppressing effects of cardiac glycosides in GSC and GBM cells. (**A**) Schematic diagram of the BLRR assay in patient-derived GSCs. GSCs from patients were stably transfected with BLRR, then sequentially transfected with lentiviruses encoding trGluc and I-SceI, followed by treatment with small-molecule candidate HDR inhibitors to assess DSB repair by BLRR assay. (**B**) The BLRR assay reveals dose-independent HDR inhibition by HDR inhibitor CAY since the Gluc signal decreases as the amount of CAY is increased, while the Vluc signal remains stable. (**C**) The BLRR ratio demonstrates HDR inhibition in GSCs by CAY. (**D, E**) Both the BLRR signal (**D**) and the BLRR ratio (**E**) reveal dose-dependent inhibition of HDR by cardiac glycosides in U87-MG GBM cells. Lanatoside C, ouabain and digoxin were treated at the indicated concentrations. (**F, G**) Both the BLRR signal (**F**) and the BLRR ratio (**G**) reveal dose-dependent HDR inhibition in GSCs treated with lanatoside C, ouabain and digoxin. BLRR signals were normalised against cell viability and results are shown as the fold change relative to DMSO-treated controls (mean ± SEM of three biological replicates). The BLRR ratio is shown as the fold change relative to DMSO (mean ± SEM of three biological replicates). (**H**) Western blot analysis revealing cardiac glycoside-induced downregulation of RAD51 in U87-MG, as well as in GSCs (**I**). RAD51 protein levels in U87-MG and GSCs treated with the indicated cardiac glycosides at 250 and 1000 nM for 24 h. Significance was calculated using one-way ANOVA as indicated, followed by Tukey’s post-hoc test *(*p* <0.05, *\*\*p* <0.01, *\*\*\*p* <0.001, *\*\*\*\*p* <0.0001).

We recently reported that pharmacological inhibition of stearoyl-CoA desaturase 1 (SCD1) with CAY10566 (CAY) downregulates the HDR protein RAD51 in GSCs as an anticancer strategy(49). Therefore, we first examined whether treating GSCs with CAY impairs HDR function. Notably, applying CAY to GSCs expressing BLRR+trGluc+I-SceI at sub-toxic nanomolar concentrations revealed a significant reduction in Gluc activity and BLRR ratio as the amount of applied CAY increased, thereby indicating an HDR-suppressing effect for CAY in GSCs (**Figure 7B, C**). Meanwhile, Vluc activity remained similar between CAY-treated and dimethyl sulphoxide (DMSO) controls. These results suggest that the BLRR accurately reports the effects of compounds on DNA DSB repair in GSCs.

We previously identified cardiac glycosides as potential glioma therapeutics, but their involvement in DSB repair remains poorly understood(50,51). To investigate the possible DSB repair-modulating effects of cardiac glycosides, human U87 GBMs as well as GSCs stably expressing BLRR were treated with low nanomolar doses of ouabain, lanatoside C, or digoxin, and BLRR assays were performed. Remarkably, cardiac glycosides significantly reduced Gluc activity and the BLRR ratio, while Vluc activity remained similar in both U87 and GSC cells, demonstrating suppression of HDR in both cell types (**Figure 7D-G**). To elucidate the mechanism of cardiac glycoside-mediated HDR inhibition, we examined RAD51 expression in treated cells, and discovered that all three cardiac glycosides triggered a dose-dependent downregulation of RAD51 protein expression, thus corroborating the decrease in HDR observed by BLRR assay (**Figure 7H, I**). These findings reinforce the antineoplastic properties of cardiac glycosides, and unveil a novel HDR-suppressing function of these natural compounds as modulators of DDR in tumour and tumour stem-like cells.

## DISCUSSION

Analysis of DNA repair is critical for the development of genome-editing tools and studying DDR in relation to (patho)physiological conditions. For instance, enhancing HDR can increase genome editing efficacy, while HDR inhibition can sensitise cancer cells to DNA-damaging anti-tumour therapies. Regarding genome editing, enhancing HDR repair pathways can improve gene knockin and knockout efficiencies during S and G2 phases since NHEJ occurs in M, G1 and G0 phases(52,53). One of the current conundrums in gene therapy is the low editing efficiency in HDR because the cell cycle is arrested in post-mitotic cells(3). However, studying DNA repair events with conventional methods such as T7E1 and Sanger sequencing is time-consuming and laborious, often requiring disruption of cells for genomic DNA extraction followed by PCR amplification and sequencing analysis(27,54). To bypass these limitations, we developed the BLRR system for the non-invasive, rapid and quantitative analysis of HDR and NHEJ repair events. Moreover, since Gluc and Vluc use different substrates, BLRR signals can be measured using the same sample, which increases the read output efficiency when screening DSB repair outcomes. Previous TLR methods have used fluorescence to detect DNA DSB repair by cell dissociation followed by flow cytometry-based analysis, which is not feasible for longitudinal studies(31). By contrast, the BLRR evaluates DSB repair by sampling only a few microliters of conditioned medium or blood to generate high signal-to-noise ratio readings of DNA repair events during longitudinal monitoring with a rapid sample turnover time (*i.e.* a few seconds per sample). Furthermore, the BLRR allows cells to remain intact for downstream applications, including sequencing and proteomic analyses.

We used both I-SceI and Cas9 to create DSBs and demonstrated that the BLRR assay reports DSB repair in a time- and event-specific manner, suggesting that it can be applied to study the dynamics between genome editing tools and DSB repair mechanisms. Interestingly, we consistently found that the introduction of trGluc donor DNA increased both HDR and NHEJ activities (**Figure 1E and Figure 3E, F**). This phenomenon concurs with observations made by Richardson *et al.* in which error-prone repair outcomes, in addition to HDR, were increased when single- and doublestranded DNA were present(40,41), thereby demonstrating the function of BLRR in accurately detecting HDR and NHEJ events. gRNA design is important for improving RNA-guided endonuclease-based editing efficiency and decreasing off-targeting effects(30,55). For example, Donech *et al.* (2014) discovered a sequence preference for gRNA activity and knockout efficiency by screening 1,841 single guide RNAs(56). Herein, BLRR analysis revealed that gRNA-3 exhibited a significantly higher HDR% and NHEJ% than gRNA-1 with the two gRNAs only 30 bp apart, demonstrating that it may be used for screening optimal gRNAs for Cas9-based editing. All tested gRNAs except gRNA3 displayed similarly low Gluc activity, and TIDER analysis revealed that gRNA1 and gRNA2 yielded the highest HDR%, while gRNA5 gave the lowest HDR%. The differences between the two analyses may be attributed to fewer HDR events, below the optimal detection limit of the assays. Meanwhile, the BLRR results also demonstrated that the closer the distance between the DSB site and the HDR arm, the higher the HDR efficiency, thereby corroborating previous findings(57). For instance, gRNA2 and gRNA4 have cut sites farther from the HDR arm than gRNA3, and both gRNA2 and gRNA4 yielded a high Vluc signal but minimal Gluc activity. Therefore, the BLRR assay is sufficiently sensitive and versatile to investigate the relationship between gRNA, DSBs and DNA repair. For example, the BLRR reporter cassette in a lentiviral vector can be cloned with HDR regions of interest to generate reporter cell lines for gRNA screening(58). The BLRR assay was able to identify ~1% of HDR and ~15% of NHEJ events in cells, and the results were highly correlated (R^2^ = >0.9) with those of NGS analysis. We further demonstrated that the BLRR enables longitudinal tracking of DSB repair events for up to 60 h. Moreover, we found that the Vluc signal declined in cells transfected with pX330-gRNA compared with the other group (**Figure 4C**). Given that NHEJ events can be elevated in the presence of donor templates(40), we speculated that the amount of transfected trGluc would decrease over the course of the experiment as cells proliferate. Consequently, cells carrying less pX330-gRNA+trGluc may proliferate faster than their counterparts, thereby resulting in an increased ratio of low plasmid-containing to high plasmid-containing cells *(i.e. an* increased low NHEJ:high NHEJ cell population ratio), and consequently a decrease in Vluc signal at the latter time points. By contrast, the NHEJ activity of the pX330-gRNA group was not potentiated by the presence of trGluc donor template from the start of the experiment, hence a slower increase in Vluc signal was observed without a decline before the end of the experiment as NHEJ accumulates. Consistently, NGS analysis showed that HDR and NHEJ events decreased at 48 h (**Figure 4D, E**), in line with the increased low plasmid-containing to high plasmid-containing cell ratio. Of note, the assay exhibited a ~6 h delay in reporting significantly increased NHEJ and HDR events compared with NGS analysis, though the general trends were similar between the two assays.

The time delay of BLRR is likely a result of the time required for the translation and release of Gluc and Vluc luciferases following DSB repair. Hence, whereas the BLRR cannot facilitate real-time detection, it enables time-lapsed monitoring of the trends of HDR and NHEJ events while keeping cells intact. By taking advantage of the high signal-to-noise ratio of Gluc and Vluc activity and the secreted luciferases, we showed that the BLRR platform can be used for longitudinal and non-invasive monitoring of HDR and NHEJ *in vivo.* We speculate that the significant increase in the BLRR signal from day 21 to day 28 likely reflects Gluc/Vluc reaching a detectable level in the blood during this period. As tumours grew, BLRR luciferases were constantly secreted, and the signals could only be detected in the blood once the signal-to-noise ratio is >1. We predict that an engineered mouse model with tissuespecific activation of BLRR could be established to study precise genome editing, including targeted delivery of transgenes, editing activity, and DDR dynamics. Efforts are currently underway to evaluate the ability of the BLRR multiplex assay to predict the efficacy of HDR inhibitors in mouse orthotopic GSC brain tumour models.

By activating intrinsic DDR, cancer cells are capable of repairing DNA damage caused by cellular stress, oxidative DNA damage in the tumour environment, and genotoxic insults induced by therapy. For instance, shifting DDR towards HDR allows tumour cells to survive exposure to DNA-damaging agents(59–61). Conversely, inhibiting or downregulating HDR proteins such as RAD51 can sensitise cancer cells to genotoxic agents by preventing DSB repair, thereby suppressing tumour growth(14,62,63). Radiation therapy and chemotherapeutics such as the alkylating agent temozolomide (TMZ) induce lethal DSB. However, increased HDR repair is identified as a common feature of several malignancies such as GBM, as well as recurrent tumours(64,65). By repairing DSB, an increase in HDR contributes significantly to acquired radioresistance(7) and TMZ resistance(65). Furthermore, GSCs are more resistant to DNA damage than their non-GSC counterparts(66,67). For instance, RAD51 contributes to the resistance of GSCs to TMZ(8), and confers resistance to radiation therapy in GBMs and GSCs. To first confirm whether BLRR can detect altered DSB repair induced by small-molecule modulators, we applied NU7441 and B02 and observed dose-dependent HDR enhancing and suppressive effects, respectively. Notably, we found that when HDR was enhanced at higher NU7441 concentrations, NHEJ was reduced, suggesting an inverse correlation between HDR and NHEJ when the repair dynamic is significantly shifted. On the other hand, we observed that both HDR and NHEJ were reduced when HDR was suppressed by B02 at higher concentrations. Consistently, we observed a decrease in DNA-PKcs expression at higher B02 concentrations, which coincides with the reduced NHEJ events (**Supplementary Figure 7**). Although the presented Gluc and Vluc values were normalised against cell viability, we also speculate that the decrease in both HDR and NHEJ may be partly attributed to cell stress and/or cell death induced by high concentrations of B02(68,69). Furthermore, we found that the BLRR ratio *(i.e.* Gluc:Vluc) may prove to be a more accurate assessment of the ability of compounds to influence DNA repair mechanisms. Taken together, the results imply that the BLRR enables analysis of the altered dynamics of DSB repair induced by small-molecule modulators.

We recently showed that inhibition of fatty acid desaturation mediated by SCD1 depletes RAD51, thereby increasing DNA damage and sensitivity to TMZ in patient-derived GSCs(28). However, whether HDR efficiency is affected by inhibition of fatty acid desaturation remains unknown. In the current study, the BLRR assay revealed dose-dependent HDR reduction induced by CAY treatment, thereby validating these findings, and confirming that pharmacological inhibition of SCD1 downregulates RAD51-mediated HDR in GBMs and GSCs. To further test the potential of the BLRR as a compound screening platform for identifying modulators of DDR, we applied lanatoside C, ouabain and digoxin, and revealed the HDR-suppressing effects of cardiac glycosides *via* RAD51 downregulation in GBMs and GSCs. These compounds, especially ouabain, displayed double-digit nanomolar potency with a >70% decrease in HDR in GSCs. Given that RAD51 activity confers resistance to radiation therapy, concomitant treatment of GBM with cardiac glycosides could potentially increase radiosensitivity. In fact, several members of the cardiac glycoside family have been previously reported to increase tumour cell death following radiation therapy(70–74). With its high sensitivity and ability to longitudinally monitor HDR and NHEJ both *in vitro* and *in vivo*, the BLRR assay serves as a versatile platform for investigating DSB repair, as well as high-throughput screening to identify and optimise gRNAs and HDR modulators.

## DATA AVAILABILITY

All correspondence and material requests should be address to CPL.

## SUPPLEMENTARY DATA

Supplementary Data are available at NAR online.

## DECLARATIONS

### Ethical approval and consent to participate

All patient samples and animals were handled under practices and operating procedures complying with the policies of the MGH Institutional Review Boards (2005P001609; MGH Animal Welfare Assurance No.: D16-00361).

## ACKNOWLEDGEMENTS

We are grateful for Dr. Hiroaki Wakimoto for providing Primary GBM cells used in this study. We thank Dr. Bakhos Tannous for providing some of the reagents used in our study and for his valuable input. We acknowledge the MGH Vector Core for producing the viral vector supported by NIH/NINDS P30NS04776. We are grateful for Dr. Mei-Yeh Lu for consultation on NGS sequencing, and service from NGS core at BRCAS in Academia Sinica. We would like to acknowledge the service provided by the DNA Sequencing Core of the Centre for Biotechnology, National Taiwan University.

---

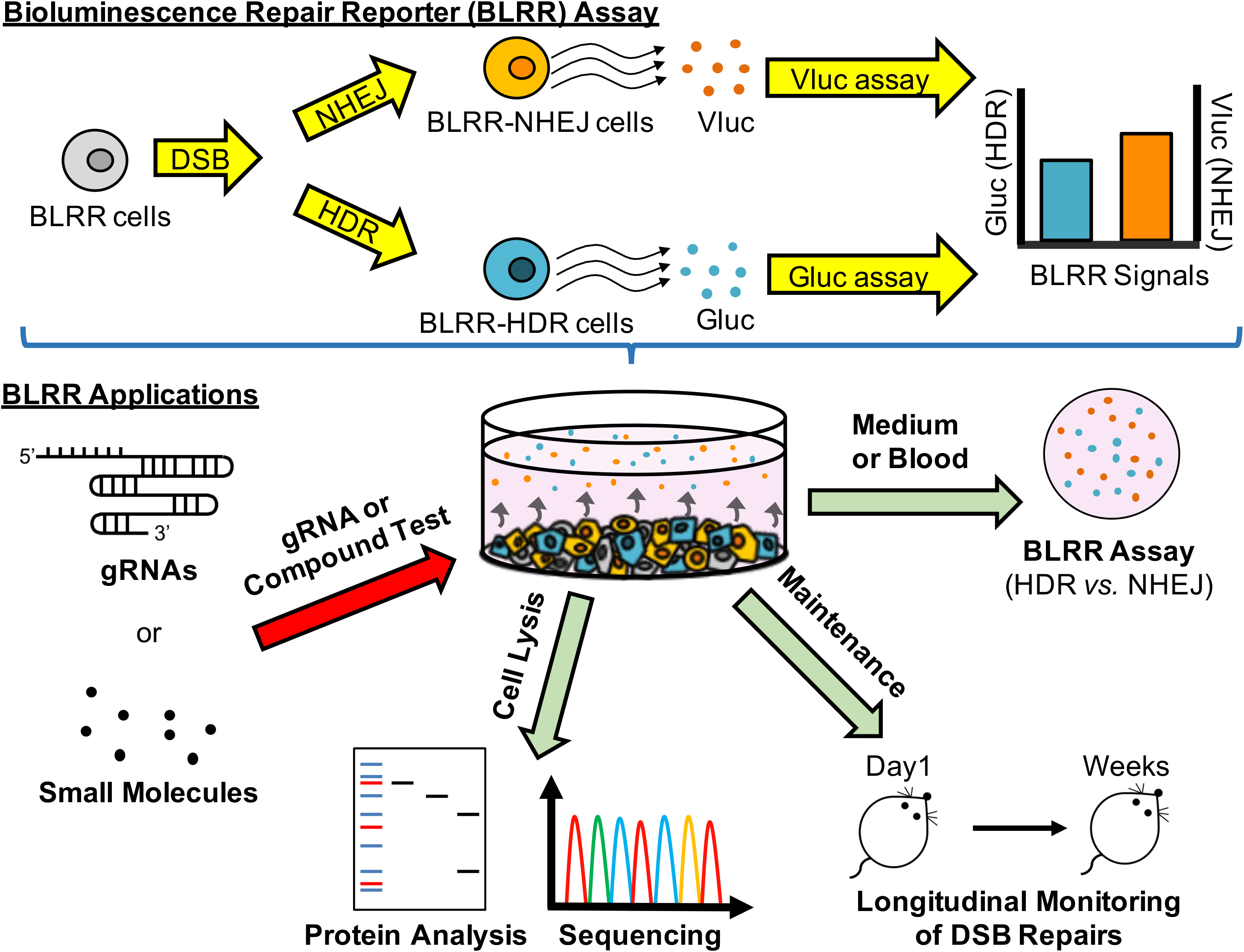

