## Supplementary_Tables_Figures for "A multiplexed bioluminescent reporter for sensitive and non-invasive tracking of DNA double strand break repair dynamics *in vitro* and *in vivo*"

**Supporting Information**

2 Supplementary Tables

8 Supplementary Figures

**Supplementary Table 1. List of primers used in this study**

| Primer Name | 5' to 3' |
|---|---|
| gRNA-1-fwd | CACCGCGACGATCGCGAGATGTTAT |
| gRNA-1 rev | AAACATAACATCTCGCGATCGTCGC |
| gRNA-2-fwd | CACCGTTATAGGTGTAGGGATAAC |
| gRNA-2-rev | AAACGTTATCCCTACACCTATAAC |
| gRNA-3-fwd | CACCGTAACAGGGTAATTCGATCTG |
| gRNA-3-rev | AAACCAGATCGAATTACCCTGTTAC |
| gRNA-4-fwd | CACCGTTATAGGTGTAGGGATAACA |
| gRNA-4-rev | AAA CTGTTATCCCTACACCTATAA C |
| gRNA-5-fwd | CACCGCGCGAGATGTTATAGGTGTA |
| gRNA-5-rev | AAA CTACACCTATAACATCTCGCGC |
| trG-fwd | TTTAGTGAACCGTCAGATCCAAGCTGGCTGCAC |
| trG-rev | ATTCGAAGCTTGAGCTCGAGCTAGTC |
| gRNA-6-fwd | CACCGTCGCGAGATGTTATAGGTGT |
| gRNA-6-rev | AAA CACACCTATAACATCTCGCGAC |
| NGS-fwd | TCGTCGGCAGCGTCAGATGTGTATAAGAGACAGGCTC AAAGAGATGGAAGCCA |
| NGS-rev | GTCTCGTGGGCTCGGAGATGTGTATAAGAGACAGCCC TTGATCTTGTCCACCTGG |
| TIDE-1-fwd | GAAGACTTCAACATCGTGGC |
| TIDE-2-Rev | TATTTTCGCAAAGTCCGTCG |
| TIDE-2-fwd | GACACCGACTCTAGTCCA |
| TIDE-2-rev | TCA CCG GAA TAA GCT AGT CG |

**Supplementary Table 2.** ***In silico*** **scores for different gRNAs**

| gRNA number | Benchling On-Target Score | CHOPCHOP Efficiency |
|---|---|---|
| 1 | 40.66 | 40.67 |
| 2 | 24.44 | 24.44 |
| 3 | 65.3 | 65.3 |
| 4 | 57.03 | 57.04 |
| 5 | 54.05 | 54.05 |
| 6 | 49.2 | 49.24 |

The insertion sequence in BLRR was analysed by Benchling (https://benchling.com) and CHOPCHOP (https://chopchop.cbu.uib.no/).

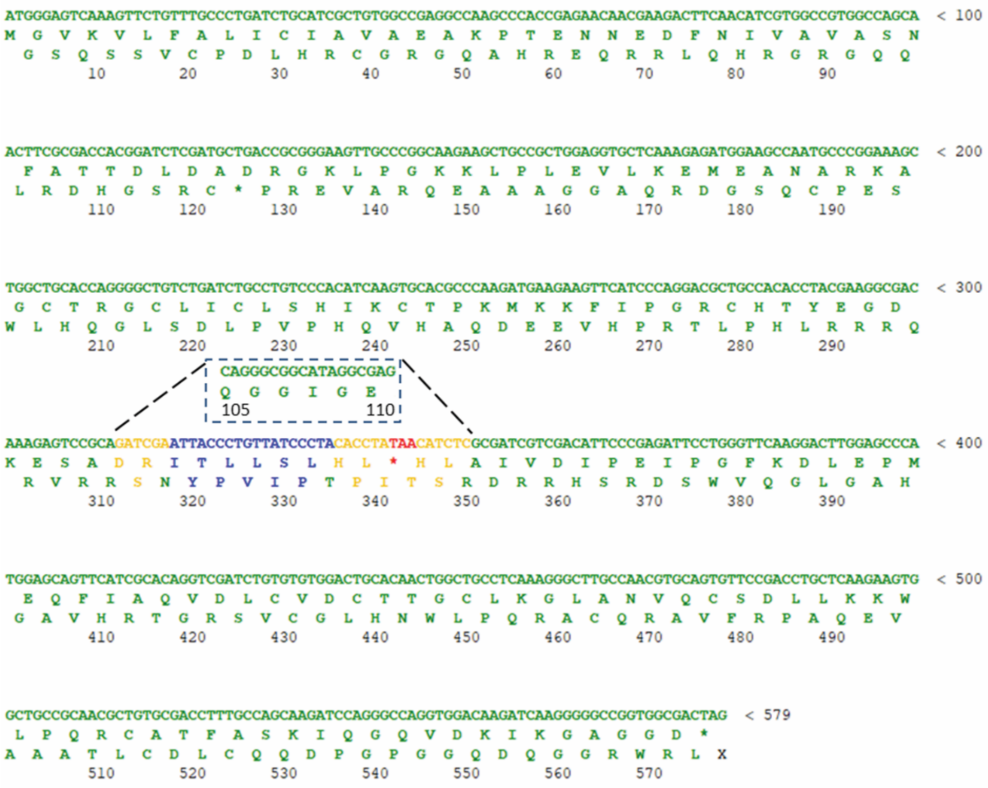

**Supplementary Figure 1. Sequence map of the silence-mutated Gluc sequence in BLRR.** Green, Gluc; yellow, spacer; red, stop codon; blue, I-SceI target site; dashed line, wild-type Gluc. Gluc, *Gaussia* luciferase

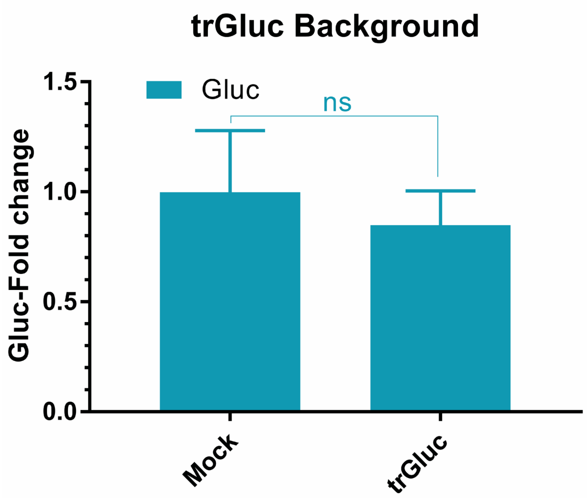


**Supplementary Figure 2. trGluc does not exhibit Gluc activity.** BLRR cells transfected with trGluc do not exhibit Gluc activity compared with mock controls. The Gluc signal was normalised against the mean value for mock controls. Data are presented as mean ± SEM of three biological replicates ($p$ >0.05; two-tailed Student t-test; ns, non-significant). Gluc, *Gaussia* luciferase; trGluc, truncated Gluc.

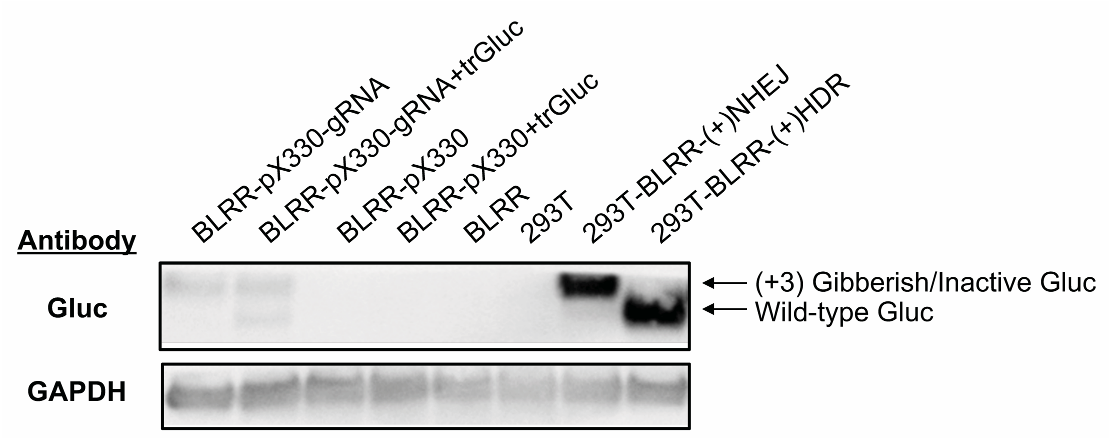


**Supplementary Figure 3. Western blotting analysis showing Gluc and T2A peptide expression following HDR and NHEJ repair events in BLRR cells.** Wild-type Gluc, 19.86 kDa; (+3) Gibberish, 23.17 kDa. BLRR cells were transfected with pX330-gRNA with or without trGluc. 293T cells transfected with BLRR-(+)NHEJ or BLRR-(+)HDR served as controls for NHEJ (*i.e.* (+3) gibberish Gluc and HDR (*i.e.* wild-type Gluc), respectively. BLRR-pX330, BLRR-pX330+trGluc, BLRR and wild-type 293T (293T) served as negative controls. GAPDH was immunoprobed as a loading control. BLRR, bioluminescent; trGluc, truncated Gluc.

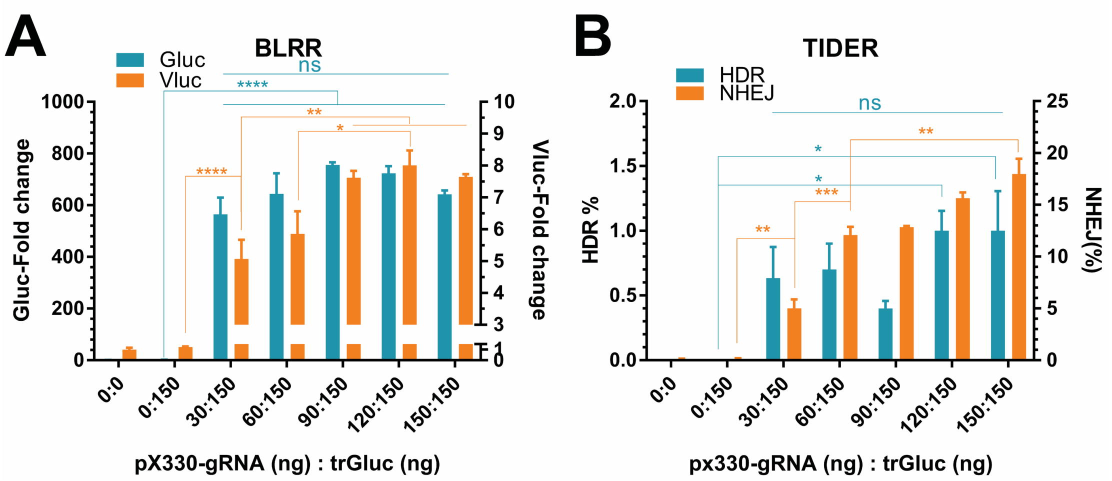


**Supplementary Figure 4. BLRR and TIDER analyses of BLRR cells transfected with increasing amounts of pX330-gRNA reveals an increase in NHEJ events.** (**A**) BLRR analysis reveals a pX330-gRNA dose-dependent increase in Vluc while elevated Gluc remains unchanged. BLRR cells were transfected with a fixed amount of trGluc and increasing quantities of pX330-gRNA. (**B**) TIDER analysis of the BLRR cells from (A) reveals a dose-dependent increase in NHEJ events. Data are presented as mean ± SEM of three biological replicates. Significance was calculated by one-way ANOVA followed by Tukey's post-hoc test (*$p$ <0.05, **$p$ <0.01, ***$p$ <0.001, **** $p$ <0.0001).

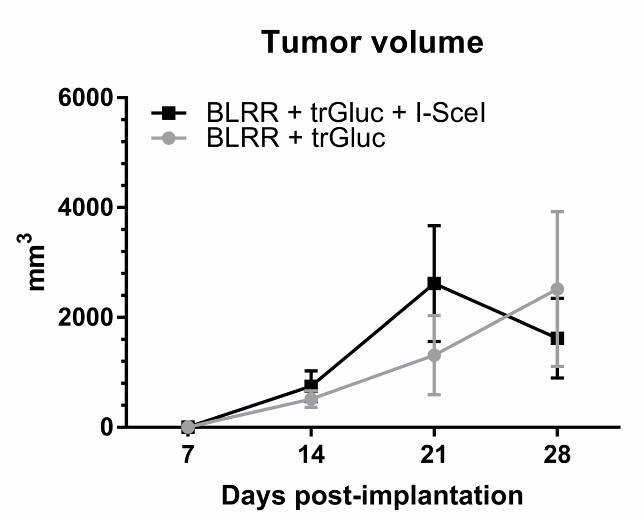


**Supplementary Figure 5. Sizes of implanted tumours over time.** Tumour growth was monitored over time by calliper measurement. Data are presented as mean ± SEM of three mice. BLRR, bioluminescent repair reporter; trGluc, truncated Gluc.

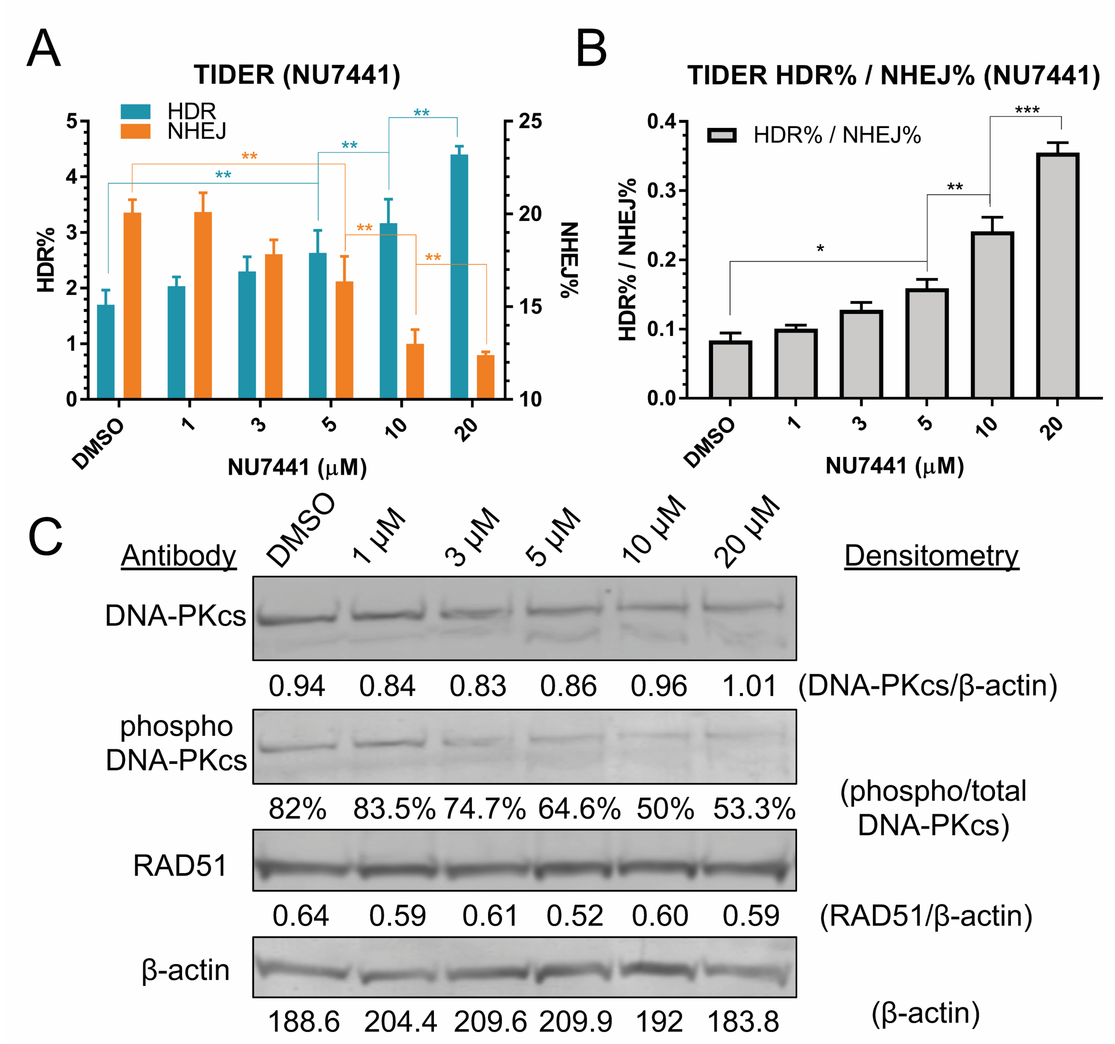


**Supplementary Figure 6. TIDER and western analysis of the HDR-enhancing effect.** (**A**) NU7441-treated BLRR cells showing a dose-dependent increase in HDR events and a decrease in NHEJ events. TIDER analysis was performed on the same cells used to generate the results shown in Figure 6A. (**B**) HDR% and NHEJ% values reveal a dose-dependent increase in HDR events. Data are presented as mean ± SEM of three biological replicates. **(C)** Western blot analysis reveals a dose-dependent decrease in the abundance of phosphorylated DNA-PKcs. Significance was calculated by one-way ANOVA compared as indicated, followed by Tukey's post-hoc test (**p* <0.05, ***p* <0.01, ****p* <0.001, *****p* <0.0001). HDR, homology-directed repair; NHEJ, non-homologous end joining; TIDER, tracking of insertions, deletions and recombination events; DNA-PKcs, DNA-dependent protein kinase catalytic subunit.

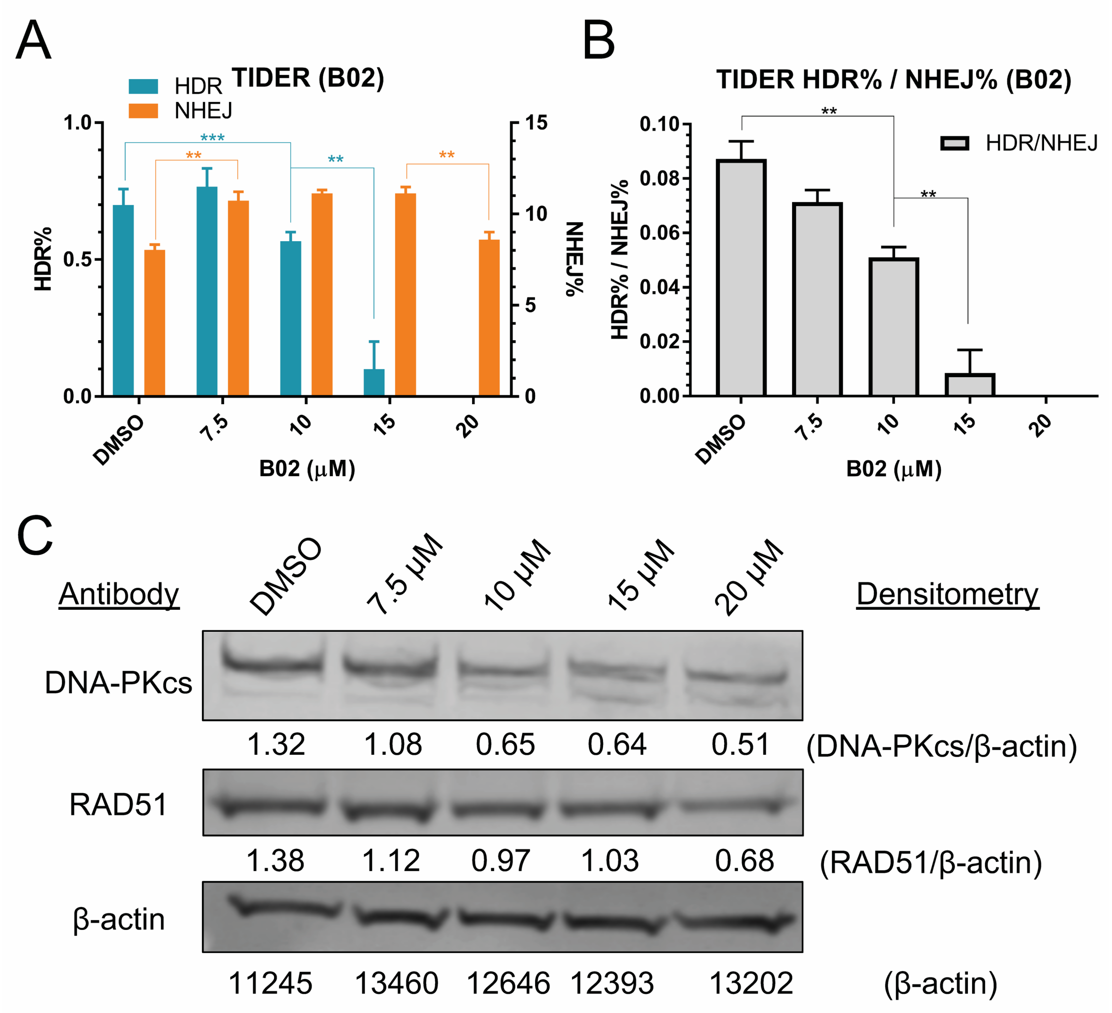


**Supplementary Figure 7. TIDER and western analysis of the HDR-inhibiting effect.** (**A**) B02-treated BLRR cells showing a dose-dependent decrease in HDR events. TIDER analysis was performed on the same cells used to generate the results shown in Figure 6D. (**B**) HDR% and NHEJ% values reveal a dose-dependent decrease in HDR events. Data are presented as mean ± SEM of three biological replicates. **(C)** Western blot analysis reveals a dose-dependent decrease in the abundance of RAD51 and DNA-PKcs. Significance was calculated using one-way ANOVA compared as indicated, followed by Tukey's post-hoc test (**p* <0.05, ***p* <0.01, ****p* <0.001, *****p* <0.0001). HDR, homology-directed repair; NHEJ, non-homologous end joining; TIDER, tracking of insertions, deletions and recombination events; DNA-PKcs, DNA-dependent protein kinase catalytic subunit.

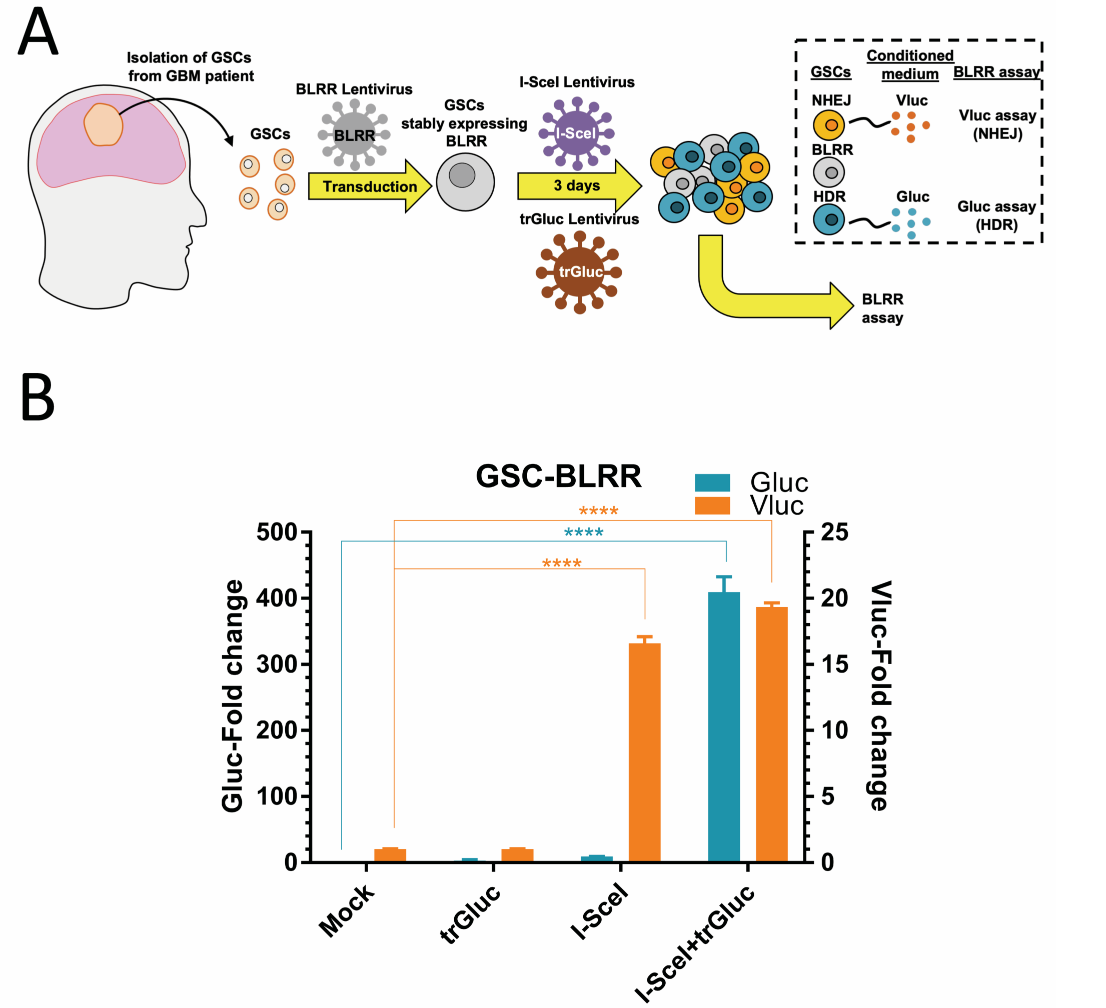


**Supplementary Figure 8. BLRR reporter controls in patient-derived GSCs.** (**A**) Schematic diagram of the BLRR assay in patient-derived GSCs. GSCs from patients were stably transfected with BLRR, then sequentially transfected with lentiviruses encoding trGluc and I-SceI, then assessed by BLRR assay. (**B**) BLRR assays demonstrate DSB repair induced by I-SceI in patient-derived GSCs. I-SceI+trGluc yields a significant increase in Gluc and Vluc signals, whereas I-SceI only yields an increase in Vluc activity, revealing BLRR specificity for reporting HDR and NHEJ events in GSCs. Mock and trGluc groups served as negative controls. Data were normalised against mock controls and results are presented as mean ± SEM of three biological replicates. Significance was calculated using one-way ANOVA compared to mock controls, followed by Tukey's post-hoc test (****$p$ <0.0001). BLRR, bioluminescent repair reporter; GBM, glioblastomas; Gluc, *Gaussia* luciferase; GSCs, GBM cancer stem cells; HDR, homology-directed repair; trGluc, truncated Gluc; Vluc, *Vargula* luciferase.
